# CaMKIIα holoenzymes self-organize into worm-like mesoscale clusters

**DOI:** 10.64898/2026.01.21.700754

**Authors:** Taisei Suzuki, Takashi Sumikama, Keisuke Matsushima, Kodai Hasegawa, Ayumi Sumino, Kenichi Umeda, Noriyuki Kodera, Tamoghna Das, Carsten Beta, Hideji Murakoshi, Mikihiro Shibata

## Abstract

Ca^2+^/calmodulin-dependent protein kinase II (CaMKII) is highly enriched in dendritic spines at concentrations comparable to those of cytoskeletal proteins and plays a central role in synaptic plasticity. During long-term potentiation (LTP), CaMKIIα further accumulates in spines. However, the mechanisms governing its higher-order organization remain poorly understood. Here, we use high-speed atomic force microscopy to visualize inter-holoenzyme interaction of CaMKIIα at mesoscopic scales (5–500 nm). Under freely diffusible conditions, CaMKIIα holoenzymes do not form stable clusters. In contrast, when spatially confined, they assemble into worm-like chain clusters mediated by kinase-domain interactions. These clusters expand upon activation, concomitant with the dissociation of the regulatory segment. Notably, the CaMKIIα P212L mutant associated with neurodevelopmental disorders, forms extensive clusters even in the basal state. Together, our findings demonstrate that CaMKIIα-CaMKIIα interactions drive mesoscale cluster formation and that precise regulation of cluster size and activation-dependent growth might be critical for synaptic signaling.

## Introduction

CaMKII is a serine/threonine kinase essential for the induction of LTP, the cellular basis of learning and memory^1–6^. Among the four CaMKII isoforms (α, β, γ, and δ), CaMKIIα and CaMKIIβ are the predominant isoforms expressed in excitatory neurons of the brain^7–10^. CaMKIIα knockout mice exhibit impaired spatial memory, illustrating its direct involvement in memory formation^11^. At the molecular level, the CaMKIIα subunit consists of a kinase domain, a regulatory segment, a linker, and a hub domain. It predominantly assembles into a dodecameric double-ring holoenzyme (Supplementary Fig. 1a, b)^12,13^. The regulatory segment contains a Ca^2+^/calmodulin (CaM)-binding site. Upon Ca^2+^/CaM binding, this segment dissociates from the kinase domain, unveiling the catalytic site and inducing an open conformation characteristic of the activated state. This conformational change enables *trans*-autophosphorylation at Thr286 (T286) within the holoenzyme, which stabilizes the kinase in its open, active configuration. Notably, phosphorylation at T286 (pT286) confers Ca^2+^-independent activity, allowing CaMKIIα to remain active after intracellular Ca^2+^ levels decline and thereby serving as a molecular mechanism for synaptic memory (Supplementary Fig. 1c).

CaMKIIα is highly expressed in excitatory neurons, accounting for approximately 2% of total protein in the hippocampus^4^, a level comparable to that of the cytoskeletal protein actin. At the subcellular level, CaMKIIα is present at high concentrations within dendritic spines, with an estimated density of approximately 960 monomers per 0.1 µm^2^ (refs.^4,5,9,14,15^). This high local concentration suggests that CaMKIIα may serve not only as a kinase but also as a structural component at synapses, potentially functioning as a scaffold in addition to its well-established enzymatic roles^16–19^. During LTP induced by high-frequency electrical stimulation, Ca^2+^ influx through NMDA receptors promotes the accumulation of CaMKIIα at the postsynaptic density (PSD) and its binding to the C-terminal region of the GluN2B subunit (GluN2B_Cter_), leading to the immobilization of CaMKIIα at synaptic sites^19–24^. This interaction is considered essential for LTP induction^19,21,25,26^. However, because the total number of CaMKIIα molecules far exceeds that of GluN2B subunits, only a fraction of CaMKIIα can directly bind GluN2B_Cter_^5,10,14,27–29^. This discrepancy suggests that CaMKIIα self-clustering is likely required to maintain its exceptionally high local concentration at activated synapses^5,6^.

Previous studies have shown that CaMKIIα clusters form both *in vitro* and in neurons upon Ca^2+^/CaM activation^30–33^. Negative-stain electron microscopy (EM) revealed small clusters or pairs consisting of two to three CaMKIIα holoenzymes in the basal state^34^. Complementary *in vitro* studies further demonstrated that CaMKIIα undergoes liquid-liquid phase separation (LLPS) in the presence of Ca^2+^/CaM and GluN2B_Cter_^35,36^. Thus, the high abundance of CaMKII within the PSD, together with its capacity for high-order assembly and LLPS, suggests that CaMKIIα clustering may contribute to the molecular mechanism underlying LTP induction^35^. However, direct visualization of CaMKIIα cluster dynamics at nanometer resolution in a high-density environment has remained elusive.

We previously employed high-speed atomic force microscopy (HS-AFM) to investigate the dynamics of CaMKIIα, CaMKIIβ, and CaMKIIα/β holoenzymes, enabling single-molecule visualization of state-dependent structural rearrangement in the CaMKII holoenzyme^13,37^. HS-AFM is a distinct technique that enables direct visualization of protein behavior in solution with nanometer resolution^37–41^.

In this study, we employed HS-AFM to observe CaMKIIα holoenzymes at concentrations comparable to those in neurons, allowing visualization of CaMKIIα cluster dynamics over length scales of several hundred nanometers. We found that CaMKIIα holoenzymes assemble into well-ordered worm-like chain clusters mediated by kinase domain interactions. Upon activation by Ca^2+^/CaM binding, these clusters expanded beyond their basal configuration. Remarkably, the *de novo* mutant CaMKIIα P212L, associated with intellectual disability (ID), formed large clusters even in the basal state. In contrast, the CaMKIIα E183V mutant, associated with autism spectrum disorder (ASD), failed to form worm-like chain clusters. These results suggest that CaMKIIα clustering may play an important role in supporting effective synaptic signalling.

## Results

### CaMKIIα holoenzymes initiate cluster formation at lower densities than within the PSD

To examine the molecular assembly patterns of CaMKIIα holoenzyme clusters at the single-molecule level, we overexpressed CaMKIIα in HEK293 cells and subsequently isolated and purified the protein using two independent affinity tags (i.e., His and Strep tags)^37^(See “Methods” for details). We first investigated whether CaMKIIα holoenzymes in solution exist as a single holoenzyme or clustered form. To address this, CaMKIIα was added directly to the imaging buffer during HS-AFM scanning, and individual holoenzymes were observed at the moment of adsorption onto the AFM substrate, allowing us to infer their cluster state in solution before surface binding (Condition 1 in Methods, Fig. 1a). Quantitative analysis indicated that more than 95% of CaMKIIα holoenzymes adsorb as single particles onto the AFM substrate in the basal state (Fig. 1b, c, and Supplementary Movie 1). We next examined the activated state by pre-incubating CaMKIIα with Ca^2+^/CaM in solution before HS-AFM observations (Condition 1 in Methods). Under these conditions, CaMKIIα holoenzymes similarly adsorbed predominantly as single holoenzymes, with more than 95% of CaMKIIα holoenzymes adsorbed as single particles onto the AFM substrate (Fig. 1c, d, and Supplementary Movie 2). Consistent with previous reports showing that CaMKIIα does not undergo LLPS in vitro, even at concentrations up to 10 µM in the absence of GluN2B ^35^, our observations indicate that CaMKIIα holoenzymes do not spontaneously assemble into higher-order clusters in solution, where they can freely diffuse.

**Fig. 1:**
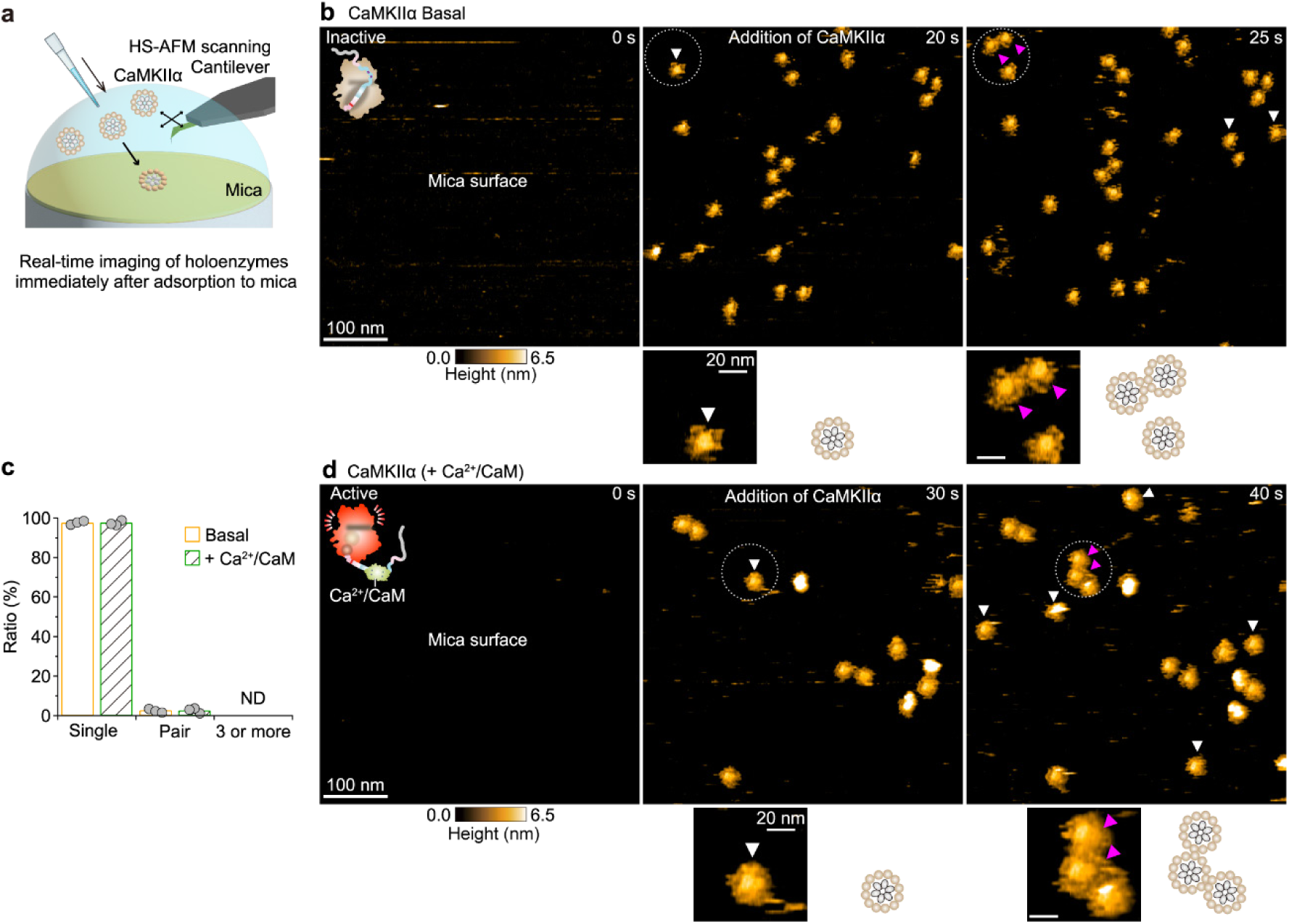
CaMKIIα holoenzymes exist predominantly as single holoenzymes in solution. **a,** Schematic illustration of the HS-AFM experimental setup for observing CaMKIIα adsorption onto the mica surface. CaMKIIα was added to the imaging buffer while continuously scanning the mica surface. **b,d** Sequential HS-AFM images of CaMKIIα adsorption onto mica in the basal(**b**; Supplementary Movie 1) and in the Ca^2+^/CaM-bound states (**d**; Supplementary Movie 2). White and magenta arrowheads indicate individual CaMKIIα holoenzymes that adsorbed on the mica surface with the corresponding frame as single or double holoenzymes, respectively. Enlarged views of the regions outlined by white dotted lines, along with the corresponding putative CaMKIIα structures, are shown below each HS-AFM image. Frame rate, 0.2 frames/s (fps). **c**, Proportion of CaMKIIα holoenzyme configuration patterns upon adsorption to the mica surface. A total of three experiments: *n* = 351 holoenzymes (basal state) and 444 holoenzymes (Ca^2+^/CaM-bound state). ND, not detected.

CaMKIIα exists at a density of 960 monomers (∼80 holoenzymes) per 0.1 μm^2^ in the PSD of excitatory neurons^4^, corresponding to ∼200 holoenzymes per 500 × 500 nm^2^. In the basal state, CaMKII binds to F-actin^16,17,42–44^, whereas upon activation, it binds to GluN2B_Cter_^43,45–49^. This spatial distribution and activity-dependent binding suggest that CaMKII diffusion is constrained within densely packed PSDs. Based on this physiological context, we sought to examine the dynamics of CaMKIIα holoenzymes under conditions that mimic both the high molecular density and constrained movement. To this end, CaMKIIα holoenzymes were added to the HS-AFM imaging buffer, and observation conditions were optimized to allow a gradual increase in the number of holoenzymes adsorbed onto the AFM substrate over time. Upon reaching a certain density state, HS-AFM observation was conducted for approximately 5 min, over a 500 × 500 nm^2^ imaging area with a temporal resolution of 5 s/frame (Condition 1 in Methods, Fig. 2a, and Supplementary Fig. 2a). At low surface densities, CaMKIIα holoenzymes diffused freely across the AFM substrate. However, as the number of CaMKIIα holoenzymes increased, inter-holoenzyme contacts became more frequent, leading to the formation of transient holoenzyme pairs and clusters (white boxes in 4). Notably, these interactions were highly dynamic, with holoenzyme pairs and clusters repeatedly forming and dissociating over time.

**Fig. 2:**
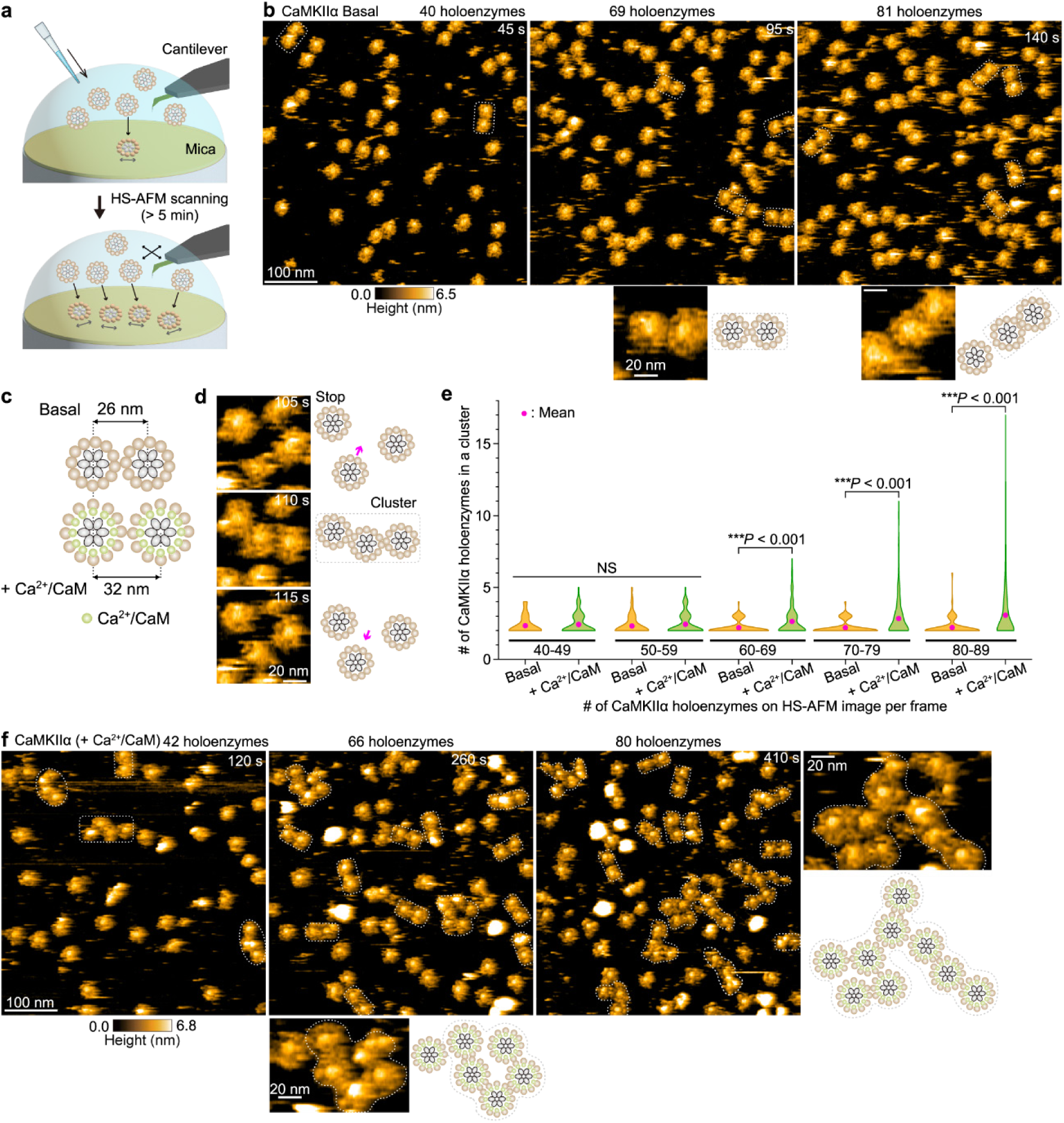
CaMKIIα holoenzymes exhibit intermolecular interactions at sub-PSD density. **a,** Schematic illustration of the HS-AFM experimental setup for observing CaMKIIα cluster formation. **b,f,** Sequential HS-AFM images showing the increasing number of CaMKIIα holoenzymes on the mica surface in the basal (**b**; Supplementary Movie 3) and the Ca^2+^/CaM-bound states (**f**; Supplementary Movie 4). White dotted lines indicate CaMKIIα clusters. Enlarged views of the regions outlined by white dotted lines, along with the corresponding putative CaMKIIα structures, are shown below each HS-AFM image. Frame rate, 0.2 fps. **c**, Schematic diagram for identifying CaMKIIα clusters (see Methods). **d**, Sequential HS-AFM images of the process of CaMKIIα cluster formation. Corresponding putative CaMKIIα structures are shown to the right of each HS-AFM image. **e**, Distribution of the number of CaMKIIα holoenzymes per cluster in the basal and Ca^2+^/CaM-bound states. HS-AFM experiments were independently repeated at least three times. Pairwise comparisons between the basal and Ca^2+^/CaM-bound states at each condition were performed using the Brunner-Munzel test. All tests are two-tailed. NS, not significant (*P* > 0.05); \*\*\**P* < 0.001. *n* = 36, 49, 77, 120, 172 clusters (basal state) and 93, 160, 390, 464, 967 clusters (Ca^2+^/CaM-bound state) for each condition.

Based on prior single-molecule observations using HS-AFM^37^, we characterized CaMKIIα cluster formation by measuring distances between holoenzymes and analyzed their behavior (Fig. 2c; see the Methods). By focusing on the initiation of clustering, we observed that at least one CaMKIIα holoenzyme stops diffusing on the substrate and contacts another holoenzyme, thereby triggering cluster formation (Fig. 2d and Supplementary Movies 3, 4). To further quantify holoenzyme mobility, we calculated the mean square displacement (MSD) by tracking the trajectories of freely diffusing CaMKIIα holoenzymes and those engaged in clusters. The result indicated that holoenzymes involved in clusters exhibit restricted diffusion (Supplementary Fig. 2b-f; see the Methods). We next analyzed the number of holoenzymes within individual CaMKIIα clusters. Under basal conditions, the cluster size remained limited to two or three holoenzymes, regardless of the total number of holoenzymes adsorbed on the substrate (Fig. 2e). In contrast, for Ca^2+^/CaM-bound CaMKIIα, when the holoenzyme number within the 500 × 500 nm^2^ area increased to 60–90 (corresponding to approximately 30– 45% of the CaMKIIα density reported in the PSD), the number of holoenzymes per cluster significantly increased compared with the basal state (Fig. 2e). Together, these results indicate that, in the activated state induced by Ca^2+^/CaM binding, high-density environments and restricted movement promote intermolecular interactions among CaMKIIα holoenzymes, leading to the formation of larger clusters.

### CaMKIIα holoenzymes form worm-like chain clusters at approximately half the density in the PSD

Under experimental conditions in which CaMKIIα was continuously supplied in the imaging buffer, individual holoenzymes diffused freely across the AFM substrate, making it difficult to resolve detailed assembly patterns of clusters at high spatial resolution. To promote stable intermolecular interactions, CaMKIIα holoenzymes were deposited onto the AFM substrate at approximately half the reported density within the PSD (30-50 holoenzymes per 300 × 300 nm^2^, corresponding to 42–69% of the PSD density). After incubation for 5 min at room temperature to allow diffusion to reach a steady-state, HS-AFM imaging was performed with a temporal resolution of 3 s/frame (Condition 2 in Methods, Fig. 3a). Under these conditions, we observed the formation of well-ordered worm-like chain clusters consisting of 2–10 CaMKIIα holoenzymes (a representative example is indicated by a white dotted line in Fig. 3b). These clusters remained stable on the AFM substrate and were basically invariant until the end of observations (Supplementary Movie 5). Further analysis of the contact points between CaMKII holoenzymes indicates that their interactions are mediated through their kinase domains. (Condition 3 in Methods, Fig. 3b).

**Fig. 3:**
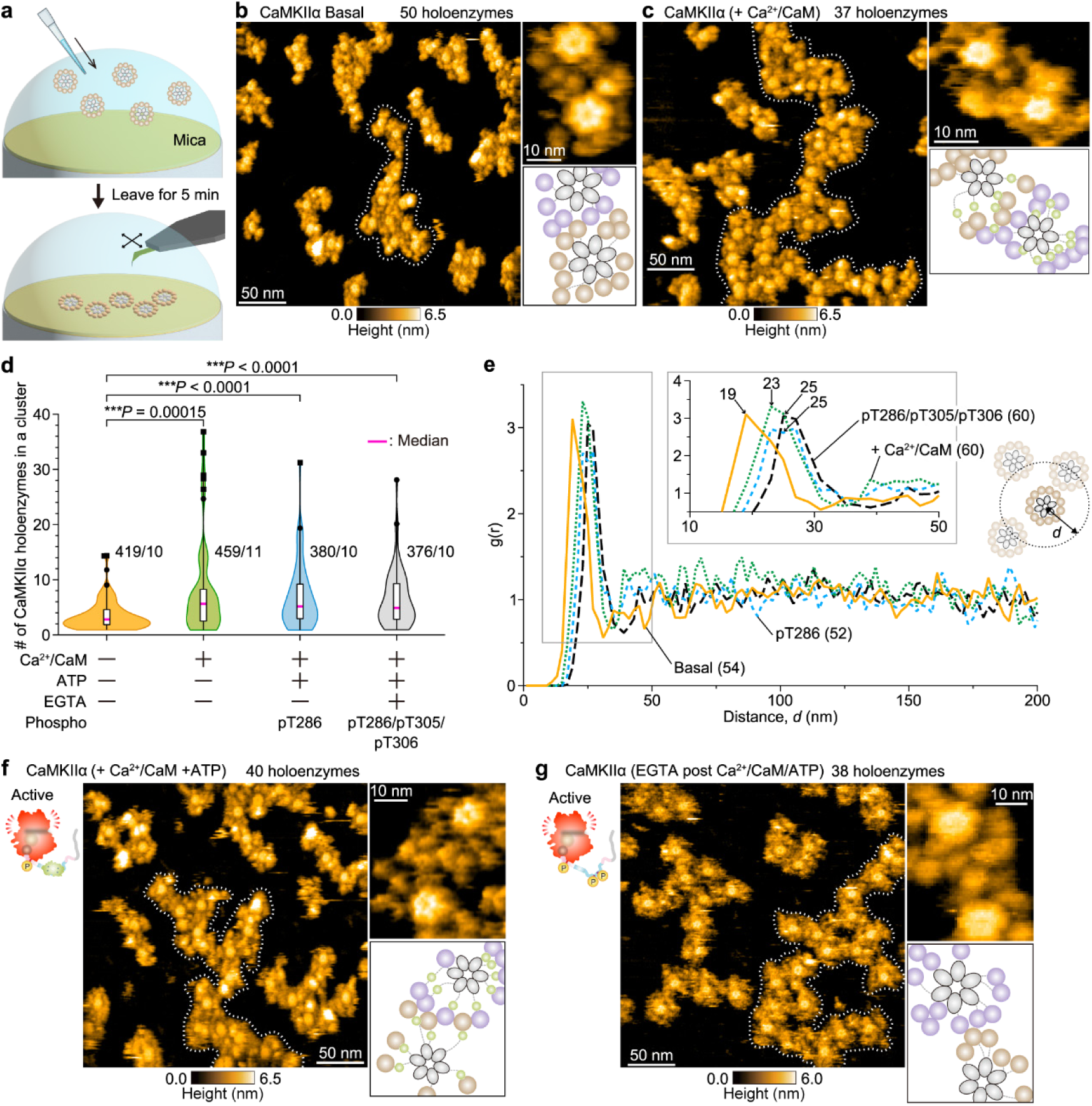
CaMKIIα holoenzymes form worm-like chain clusters via kinase-domain interactions, and these clusters expand upon Ca^2+^/CaM binding. **a,** Schematic illustration of the HS-AFM experimental setup for observing stable CaMKIIα clusters. CaMKIIα (300 nM) was deposited onto a mica surface and incubated for 5 min before HS-AFM scanning. **b,c,f,g,** HS-AFM images of stable CaMKIIα clusters in the basal state (**b**; Supplementary Movie 5), Ca^2+^/CaM-bound state (**c**; 1 mM CaCl_2_, 300 nM CaM; Supplementary Movie 6), pT286 (**f**; 1 mM CaCl_2_, 300 nM CaM, and 1 mM ATP; Supplementary Movie 7), and pT286/pT305/pT306 (**g**; Supplementary Movie 8). White dotted lines indicate representative CaMKIIα clusters. Enlarged HS-AFM images highlighting intermolecular interactions of CaMKIIα, along with the corresponding putative CaMKIIα structures (right). Frame rate, 0.33 fps. **d**, The number of CaMKIIα holoenzymes per cluster under different experimental conditions. Sample sizes (holoenzymes/images) are indicated in the figure. In box plots, the center line represents the median, box limits delineate 25^th^ and 75^th^ percentiles (IQR), and whiskers extend to the minimum and maximum values. Outliers are indicated with black markers. Kruskal‒Wallis test with Dunn’s post hoc test. All tests are two-tailed. NS, not significant (*P* > 0.05); \*\*\**P* < 0.001. **e**, Radial distribution function (RDF; Supplementary Fig. 3, and Methods) calculated from the spatial coordinates of the hub assembly of CaMKIIα holoenzymes under different conditions. The number of analyzed images is indicated in the figure. A schematic illustration of the RDF and an enlarged view of the region outlined by the gray line are shown. Bin width, 2 nm.

Next, we examined how structural changes in CaMKIIα induced by Ca^2+^/CaM binding influence the formation of worm-like chain clusters. In this experiment, CaMKIIα was pre-incubated with Ca^2+^/CaM in a tube and subsequently applied onto the AFM substrate. Similar to the basal state, the holoenzyme density was adjusted to 30–50 particles per 300 × 300 nm^2^ (Condition 2 in Methods). As expected, under these conditions, worm-like chain clusters were observed in Ca^2+^/CaM-bound CaMKIIα, with intermolecular interactions mediated through the kinase domains (white dotted line in Fig. 3c and Supplementary Movie 6). However, quantitative analysis of cluster composition (see Methods) revealed that the number of holoenzymes per cluster was significantly increased in the Ca^2+^/CaM-bound state compared with the basal state (Fig. 3d). To quantify the spatial organization of CaMKIIα holoenzymes on the AFM substrate, we determined the center coordinates of each holoenzymes (hub assembly) and calculated the radial distribution function (RDF)^50^ (See Methods). The RDF represents the probability of finding neighboring holoenzymes as a function of interparticle distance (Distance, *d*, in nm). Positions of the peaks in the RDF indicate distances where there is a high probability of neighboring holoenzymes. In the basal state, the RDF exhibited a peak at approximately 19 nm, whereas in the Ca^2+^/CaM-bound state this peak shifted outward by ∼4 nm to approximately 23 nm (Fig. 3e). Previous single-molecule imaging of CaMKIIα has shown that Ca^2+^/CaM binding causes an approximately 3 nm extension between the hub and kinase domains^37^. This coincides with the RDF peak shift, suggesting that within clustered CaMKIIα holoenzymes, the kinase domain adopts an extended structure in the radial direction.

We further evaluated cluster stability by converting the RDF to free energy profiles for each experimental condition (Methods). The analysis revealed that the pairwise free energy was nearly identical in both the basal and Ca^2+^/CaM-bound state (Supplementary Fig. 3a). In contrast, when the free energy was calculated on a per-holoenzyme basis within each cluster (Methods), we found that the basal state exhibited a free energy of - 0.31 ± 0.15 kcal/mol, whereas the Ca^2+^/CaM-bound state exhibited a more favorable free energy of −0.49 ± 0.30 kcal/mol (Supplementary Fig. 3b and Table 1). These results indicate that Ca^2+^/CaM binding enhances cluster stability. Accordingly, in the steady-state where holoenzyme diffusion has stabilized, CaMKIIα holoenzymes assemble into worm-like chain clusters containing several to ten holoenzymes linked through their kinase domains. Notably, in the Ca^2+^/CaM-bound state, a large, stable cluster was observed, consisting of more than 20 CaMKIIα holoenzymes. This extensive clustering is likely driven by Ca^2+^/CaM-induced structural changes in the kinase domains, which adopt a more extended conformation, thereby promoting multivalent intermolecular interactions.

**Table 1:** Average free energy per holoenzyme within each cluster under different experimental conditions (see also Supplementary Fig. 3b; “Methods” for details).

| Condition | AVG (E/molecule) [kcal/mol] |
| --- | --- |
| CaMKIIα<sub>WT</sub> Basal | -0.31 ± 0.15 |
| CaMKIIα<sub>WT</sub> +Ca<sup>2+</sup>/CaM | -0.49 ± 0.30 |
| CaMKIIα<sub>WT</sub> pT286 | -0.33 ± 0.15 |
| CaMKIIα<sub>WT</sub> pT286/pT305/pT306 | -0.31 ± 0.18 |

### CaMKIIα holoenzymes form large clusters in autophosphorylation state

Following Ca^2+^/CaM binding, CaMKIIα undergoes its initial autophosphorylation at Thr286 (Ca^2+^/CaM+pT286) through ATP hydrolysis. Subsequently, upon Ca^2+^/CaM dissociation, additional autophosphorylation occurs at Thr305 and Thr306, resulting in the pT286/pT305/pT306 state (Supplementary Fig. 1c). To examine how these phosphorylations influence worm-like chain clustering, each autophosphorylation reaction was induced in solution, followed by HS-AFM observations. The phosphorylation states of CaMKIIα were confirmed by biochemical assays (Supplementary Fig. 4a). In both the Ca^2+^/CaM+pT286 and pT286/pT305/pT306 states, worm-like chain clusters linked through the kinase domain were observed (Fig. 3f, g, and Supplementary Movies 7, 8). Quantitative analysis revealed that the number of holoenzymes per cluster significantly increased compared with the basal state, but was not significantly different from that observed in the Ca^2+^/CaM-bound state (Fig. 3d). Consistently, the RDF exhibited a peak at approximately 25 nm for both the Ca^2+^/CaM+pT286 and pT286/pT305/pT306 states, representing an outward shift relative to the basal state and closely resembling the Ca^2+^/CaM-bound condition (Fig. 3e). Together, these results indicate that autophosphorylation stabilizes an extended kinase-domain conformation and promotes the formation of larger CaMKIIα holoenzyme clusters.

Next, to determine which activated states of CaMKIIα most strongly influence larger cluster formation, we performed HS-AFM observations using two phosphorylation-dead mutants, T286A (CaMKIIα_T286A_) and T305A/T306V (CaMKIIα_T305A/T306V_) (Condition 2 in Methods). Biochemical assays confirmed the phosphorylation states of these mutants (Supplementary Fig. 4b, c). In CaMKIIα_T286A_, where autophosphorylation is absent, large clusters were observed solely in the Ca^2+^/CaM-bound state (Figure 4a-c, g). This suggests that Ca^2+^/CaM binding alone is essential for the formation of larger clusters. In CaMKIIα_T305A/T306V_, large clusters were observed in both the Ca^2+^/CaM+pT286 and the pT286 states (following the addition of EGTA to dissociate Ca^2+^/CaM after Ca^2+^/CaM/ATP induction) (Figure 4d-f, h). These observations indicate that large cluster formation can be driven either by Ca^2+^/CaM binding or T286 phosphorylation. Importantly, both activation mechanisms share a common structural consequence, i.e, the displacement of the regulatory segment from the kinase domain, which facilitates the formation of substantially larger CaMKIIα holoenzyme clusters compared with the basal state.

**Fig. 4:**
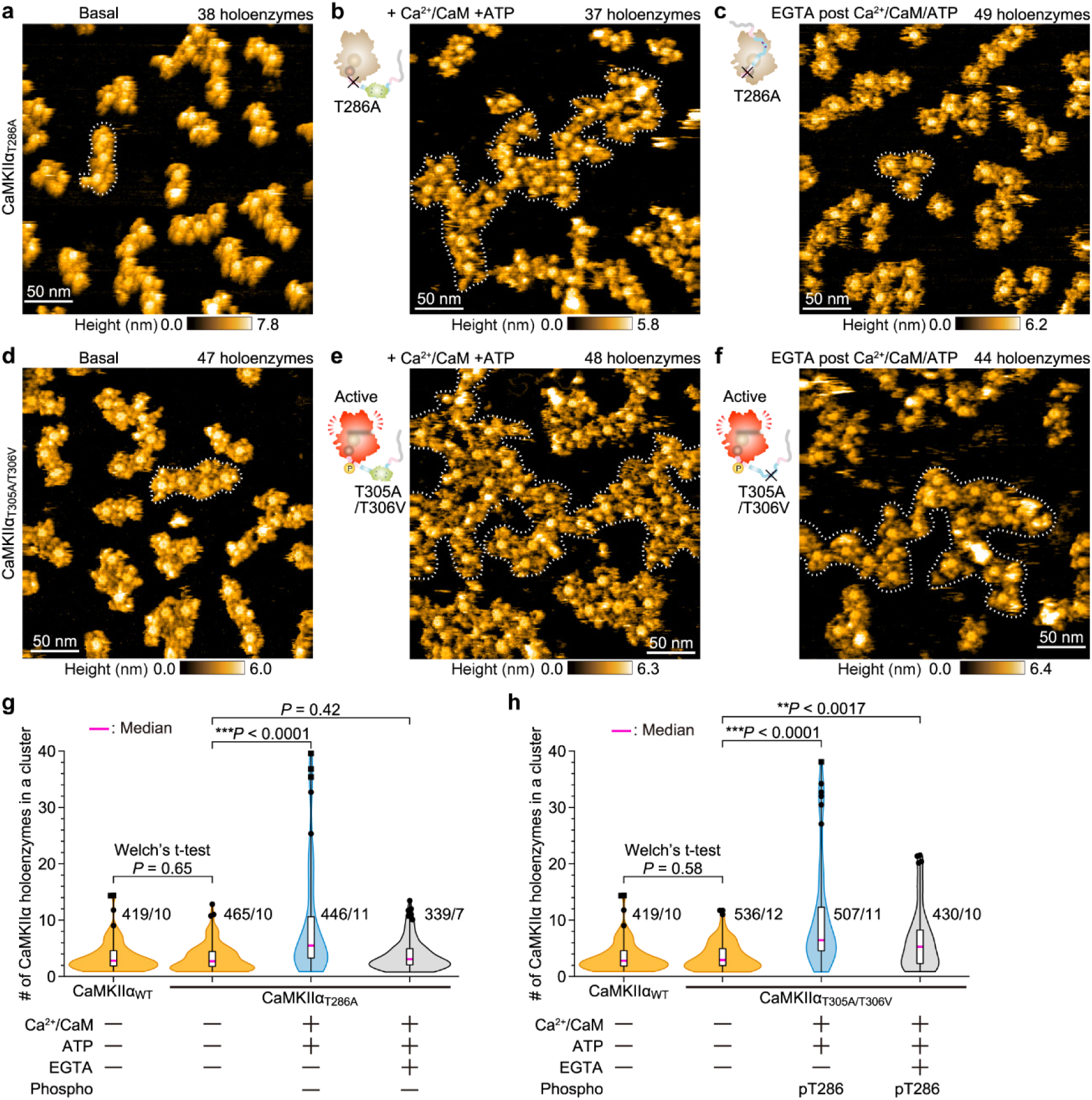
Detachment of the regulatory segment from the kinase domain promotes expansion of CaMKIIα clusters. **a–f,** Representative HS-AFM images of stable CaMKIIα_T286A_ (**a–c**) and CaMKIIα_T305A/T306V_ (**d–f**) clusters. Panels show the basal state (**a,d**), the Ca^2+^/CaM-bound state in the presence of ATP (**b,e**), and the state after CaM release by EGTA treatment (i.e., inducing pT305/pT306) (**c,f**). Frame rate, 0.33 fps. **g,h,** Quantification of holoenzymes per cluster for CaMKIIα_T286A_ (**g**) and CaMKIIα_T305A/T306V_ (**h**) under different experimental conditions. Sample sizes (holoenzymes/images) are indicated in the figure. In box plots, the center line represents the median, box limits delineate 25^th^ and 75^th^ percentiles (IQR), and whiskers extend to the minimum and maximum values. Outliers are indicated with black markers. Welch’s t-test was used to compare WT and mutant in the basal state. Comparison across conditions for each mutant was performed using the Kruskal-Wallis test with Dunn’s post hoc. All tests are two-tailed. NS, not significant (*P* > 0.05); \*\**P* < 0.01, \*\*\**P* < 0.001.

### CaMKIIα mutants associated with NDDs exhibit cluster sizes distinct from WT

To explore the potential biological implications of worm-like chain clusters formed among CaMKIIα holoenzymes, we next performed HS-AFM observations on previously reported *de novo* CaMKIIα mutants associated with neurodevelopmental disorders (NDDs). Specifically, we focused on the P212L mutant (CaMKIIα_P212L_)^51–53^, identified in patients with intellectual disability (ID), and the E183V mutant (CaMKIIα_E183V_)^51,54^, identified in patients with autism spectrum disorder (ASD). P212 lies slightly distal to the regulatory segment of the kinase domain (Fig. 5a), and heterozygous P212L knock-in mice have been reported to exhibit ID-like phenotypes^53^. In contrast, E183 resides near the outer surface of the kinase domain (Fig. 5a), and E183V knock-in mice exhibit ASD-like behavioral phenotypes, including hyperactivity, impaired social interaction, and increased repetitive behaviors^55^. Notably, since neither mutation affects the catalytic lysine (K43) nor any known phosphorylation sites, they are not expected to directly abolish kinase activity. However, given their strong association with NDDs, these variants have attracted considerable interest as potential molecular determinants of disease mechanisms^51,53,55,56^. Accordingly, we overexpressed CaMKIIα_P212L_ and CaMKIIα_E183V_ homomeric holoenzymes in HEK293 cells using a similar procedure to that for wild-type (WT) (Supplementary Fig. 5), followed by HS-AFM observations to assess their clustering behavior.

**Fig. 5:**
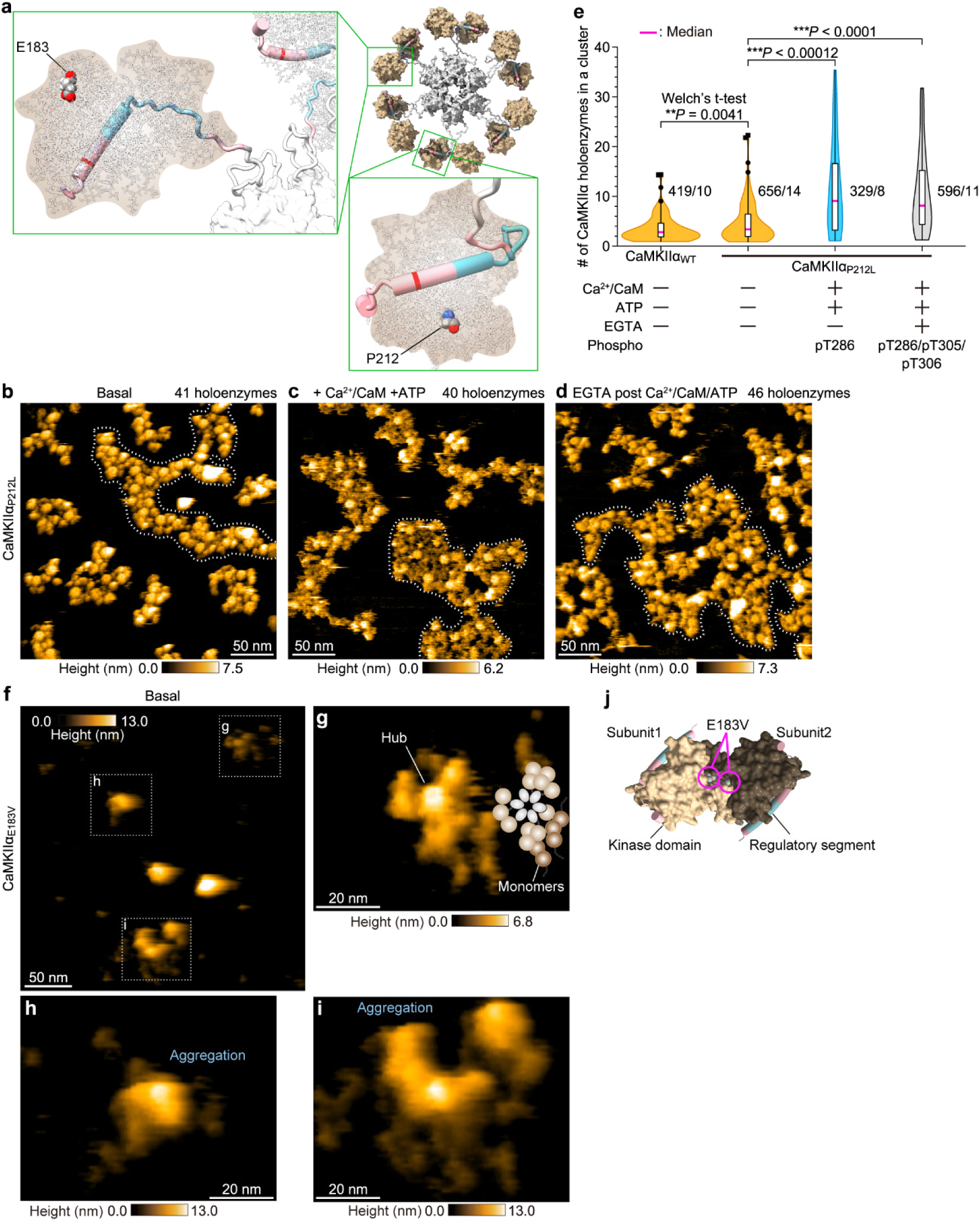
NDDs-associated CaMKIIα mutants form clusters distinct from WT. **a,** Structural locations of P212 and E183. P212 and E183 are shown as spheres in an enlarged view of a CaMKIIα subunit from the pseudoatomic EM-derived model^12^. **b–d,** Representative HS-AFM images of stable CaMKIIα_P212L_ clusters in the basal state (**b**), the Ca^2+^/CaM-bound state with ATP (**c**), and the state after CaM release by EGTA treatment (i.e., inducing pT305/pT306) (**d**). Frame rate, 0.33 fps. **e,** Quantification of holoenzymes per cluster for CaMKIIα_P212L_ under the indicated experimental conditions. Sample sizes (holoenzymes/images) are indicated in the figure. In box plots, the center line represents the median, box limits delineate 25^th^ and 75^th^ percentiles (IQR), and whiskers extend to the minimum and maximum values. Outliers are indicated with black markers. Welch’s t-test was used to compare WT and mutant in the basal state. Comparison across conditions for the mutants was performed using the Kruskal-Wallis test with Dunn’s post hoc. All tests are two-tailed. NS, not significant (*P* > 0.05); \*\**P* < 0.01, \*\*\**P* < 0.001. **f,** Representative HS-AFM images of CaMKIIα_E183V_ on a pillar[5]arene-modified mica surface. Frame rate, 0.33 fps. **g–i**, Enlarged views of regions indicated by white dotted lines in (**f**). A representative holoenzyme forming a complete dodecamer with a central hub domain is shown, although several peripheral subunits appear tangled with monomers (**g**). A holoenzyme forming an aggregate lacking dodecameric organization (**h**). A holoenzyme forming a large, disordered aggregate (**i**). Frame rate, 3.33 fps. **j**, AlphaFold3-based prediction of kinase-kinase interactions for CaMKIIα_E183V_ using residues 1–314 (kinase domain with regulatory segment). The highest scoring model is shown; a sphere indicates the E183V position (see also Table 2, and Supplementary Fig. 7).

**Table 2.** AlphaFold3 predictions for CaMKIIα_WT_ and CaMKIIα_E183V_. For each model, the interface predicted template modeling score (ipTM), the predicted template modeling score (pTM), and the “fraction disordered” are shown. ipTM assesses the accuracy of predicted interfacial interactions, and pTM evaluates the overall accuracy of the predicted structure. The “fraction disordered” indicates the proportion of residues predicted to be in a disordered conformation.

| Model | Ranking<br>score | iptm | ptm | Fraction<br>disordered |
| --- | --- | --- | --- | --- |
| **CaMKIIα<sub>WT</sub>** |  |  |  |  |
| Model 1 | 0.19 | 0.09 | 0.50 | 0.04 |
| Model 2 | 0.19 | 0.1 | 0.50 | 0.04 |
| Model 3 | 0.21 | 0.12 | 0.50 | 0.03 |
| **CaMKIIα<sub>E183V</sub>** |  |  |  |  |
| Model 1 | 0.31 | 0.24 | 0.49 | 0.03 |
| Model 2 | 0.36 | 0.29 | 0.49 | 0.05 |
| Model 3 | 0.32 | 0.25 | 0.49 | 0.04 |

Worm-like chain clusters were observed in CaMKIIα_P212L_, and both Ca^2+^/CaM+pT286 and pT286/pT305/pT306 states tended to form large clusters similar to WT (Condition 2 in Methods, Fig. 5b-d). Importantly, comparison with WT in the basal state revealed that CaMKIIα_P212L_ predominantly formed larger clusters (Fig. 5e). In contrast, HS-AFM observations of CaMKIIα_E183V_ revealed dispersed aggregates within a 300 × 300 nm^2^, without worm-like chain clusters observed in WT (Condition 3 in Methods, Fig. 5f). Although high spatiotemporal resolution imaging (<1 nm, 0.3 s/frame) demonstrated that a few holoenzymes retained the canonical 12-meric hub assembly (central gear-like structure in Fig. 5g), the majority of aggregates exhibited kinase domains arranged in disordered and entangled structures (Fig. 5h, i). Consistent with these observations, previous size-exclusion chromatography (SEC) analyses of CaMKIIα_E183V_ reported formation of large aggregates^55^. Together, these results suggest that CaMKIIα_P212L_ intrinsically favors the formation of large clusters even in the basal state, whereas CaMKIIα_E183V_ is impaired in its ability to assemble stable 12-mer holoenzyme and worm-like chain clusters. Considering the relevance to NDDs, the ability to form and maintain appropriately sized worm-like chain clusters among CaMKIIα holoenzymes suggests an important role in effective synaptic signaling and neuronal function.

## Discussion

CaMKIIα is highly enriched at excitatory postsynapses and is essential for synaptic plasticity through its high local abundance and interactions with diverse substrates^2,5,6,18^. However, how CaMKIIα holoenzymes assemble into higher-order clusters has remained unclear. Using HS-AFM, we directly visualized CaMKIIα cluster dynamics at mesoscopic scales (5–500 nm). We show that CaMKIIα forms worm-like chain clusters via kinase-domain-mediated intermolecular interactions, that Ca^2+^/CaM binding promotes an extended holoenzyme conformation and cluster expansion, and that autophosphorylation at T286 stabilizes the enlarged clusters. Moreover, NDD-associated CaMKIIα variants display abnormal cluster sizes even in the basal state (Supplementary Fig. 8a).

Our HS-AFM observations indicate that cluster formation can be initiated by locally confined CaMKIIα holoenzyme and that subsequent cluster growth requires a conformational change in CaMKIIα holoenzymes—specifically, radial extension of the kinase domains upon Ca^2+^/CaM binding. This suggests that spatial confinement and conformational expansion jointly facilitate CaMKIIα cluster assembly, independent of a specific surface context. In this framework, increasing the effective particle size of CaMKIIα holoenzymes is expected to promote cluster growth. To test this hypothesis, we performed clustering simulations based on holoenzyme trajectories obtained from HS-AFM and examined how restricting holoenzyme motion or increasing particle size affects cluster size (see Methods, Supplementary Fig. 6, Supplementary Tables 1, 2, Supplementary Discussion, and Supplementary Movies 9-12). Simulations comparing particles with radii of 13 nm (basal state) and 16 nm (Ca^2+^/CaM-bound state) revealed that the larger particles formed clusters with a significantly greater average number of holoenzymes, indicative of enhanced cluster growth (Supplementary Fig. 6h, i). Additionally, to replicate the anchoring of CaMKII to the synapse via NMDA receptors at the synapse, 13 fixed holoenzymes were introduced to represent CaMKII-NMDAR complexes, based on previous studies showing 4–18 NMDA receptors per synapse^57^. This approach led to a significant increase in the average number of holoenzymes per cluster (Supplementary Fig. 6h, i). Together, these simulations demonstrate that both an increase in the effective size of CaMKIIα holoenzymes and localized constraints promote the expansion of the CaMKIIα cluster.

Previous studies of CaMKIIα_P212L_ knock-in mice have reported increased autophosphorylation, resulting in abnormally elevated kinase activity. Consistent with this molecular phenotype, these mice exhibited hyperexcitable hippocampal LTP that can be induced by subthreshold low-frequency stimulation^56^. In silico analysis using FoldX^58^ predicts that P212L substitution destabilizes the hydrophobic core of the kinase domain and weakens interactions between the kinase domain and the regulatory segment^52^. Consequently, P212L mutations likely loosen autoinhibited conformation and bias CaMKIIα toward an extended structural state in which the regulatory segment is disengaged. Our HS-AFM observations indicate that the extended structure of the kinase domain promotes the formation of larger CaMKIIα clusters. Accordingly, CaMKIIα_P212L_ adopts a conformation closer to the extended structure in the basal state, thereby facilitating the formation of enlarged clusters (Supplementary Fig. 8a). Such enhanced basal clustering of CaMKIIα_P212L_ may lower the threshold for LTP induction by stimuli that are insufficient to elicit LTP in WT neurons, providing a potential mechanistic link between altered mesoscale organization of CaMKIIα and the hyperexcitability observed in NDDs.

CaMKIIα_E183V_ knock-in mice showed reduced CaMKIIα expression throughout the forebrain and decreased localization to synaptic subcellular fractions. In cultured neurons, the E183V mutation impairs targeting to dendritic spines. Neuronal expression of CaMKIIα_E183V_ also increases dendritic branching, accompanied by decreased dendritic spine density and reduced excitatory synaptic transmission^55^. Our HS-AFM observations further showed that CaMKIIα_E183V_ fails to assemble stable 12-meric holoenzymes and worm-like chain clusters. To assess how the E183 to Valine substitution alters kinase-kinase interactions, we predicted the dimer structure of CaMKIIα_E183V_ (residue 1-314, corresponding to the kinase domain and regulatory segment) using AlphaFold3 (AF3)^59^. Overall confidence was low for both WT and E183V predictions, but the highest-scoring E183V model displayed V183 residues in close proximity to one another (Fig. 5j, Table2, and Supplementary Fig. 7). This close-contact arrangement was not observed in WT models, implying that E183V substitution strengthens inter-kinase contacts relative to WT and thereby promotes aberrant aggregation.

Taken together, these two NDD-associated CaMKIIα variants define opposing perturbations of mesoscale CaMKIIα organization: whereas the P212L mutation biases CaMKIIα toward an extended conformation that promotes excessive worm-like cluster expansion, the E183V mutation disrupts proper holoenzyme assembly and cluster formation, instead favoring aberrant inter-kinase contacts and aggregation. This contrast highlights the existence of an optimal regime of CaMKIIα clustering, deviations from which—either excessive expansion or impaired assembly—can destabilize synaptic signaling and plasticity.

Our HS-AFM images revealed that CaMKIIα clusters are connected via the kinase domains. However, we were unable to visualize the distinct structure of the kinase domain at the linkage sites. This limitation may be due to the spatial resolution of HS-AFM or the possible absence of a distinct linkage pattern (Fig. 3). Additionally, using AF3, docking structure predictions for two CaMKIIα_WT_ subunits (kinase domain with regulatory segment) resulted in various kinase domain contact models with similar ranking scores (Table 2). These findings suggest that the stable structure formed by the contact between two kinase domains is not unique, indicating that no specific interactions are mediated. Furthermore, the free energy profile based on HS-AFM data indicated that the CaMKIIα-CaMKIIα interaction was approximately −0.7 kcal/mol (Supplementary Fig. 3a). This value is comparable to thermal fluctuations at room temperature (*k*_B_*T* = 0.6 kcal/mol), supporting the hypothesis that interactions between CaMKIIα holoenzymes are governed by non-specific interactions. The dissociation of the regulatory segment upon Ca^2+^/CaM binding alters the charge and hydrophobicity on the kinase domain surface by exposing S- and T-sites, potentially modulating CaMKIIα-CaMKIIα interactions. This modulation may contribute to cluster growth and stability, in addition to increasing holoenzyme size through Ca^2+^/CaM binding.

Previous studies using colloids have demonstrated that isolated, chain-like clusters emerge during the initial stages of gelation, suggesting that short-range intermolecular interactions and hydrodynamic interactions are essential^60^. Under the influence of hydrodynamic interactions, the solvent’s incompressibility prevents particles from converging directly toward the center, instead inducing lateral flow. This facilitates the formation of chain-like clusters. We hypothesize that CaMKIIα worm-like chain clusters exhibit a similar mechanism. As the density of CaMKIIα holoenzymes increases, non-specific interactions between holoenzymes arise, producing an effect analogous to the initial stages of a colloid gelation. This likely promotes the assembly of worm-like chain clusters of CaMKIIα holoenzymes.

Based on these results, we propose a model for CaMKIIα localization to synapse during LTP (Supplementary Fig.8b). In the basal state, a CaMKIIα holoenzymes either diffuse in spines or binds F-actin^16,17,42,44^, forming small, worm-like chain clusters of a few holoenzymes. LTP-evoked Ca^2+^ influx promotes Ca^2+^/CaM binding, which shifts CaMKIIα toward an extended conformation that permits binding to the GluN2B_Cter_ at the PSD^17,19,42,61^. This CaMKIIα-GluN2B complex acts as a nucleation site; the cluster enlarges by recruiting adjacent CaMKIIα holoenzymes via kinase-domain contacts. Recent studies have shown that Ca^2+^/CaM binding to CaMKII, followed by interaction with GluN2B_Cter_, is necessary for the early phase of spine structural plasticity during LTP^19^. Therefore, alterations in CaMKIIα-CaMKIIα interactions modulate cluster dimensions and the local abundance of CaMKIIα in the spine, thereby influencing the induction and expression of synaptic plasticity.

Our HS-AFM observations are limited to monitoring intermolecular interactions of CaMKIIα in the two-dimensional (2D) space on the AFM substrate. In addition, under current experimental conditions, both Ca^2+^/CaM and CaMKIIα exhibit non-specific adsorption to the AFM substrate, preventing real-time visualization of the sequential processes involving Ca^2+^/CaM binding to basal CaMKIIα and the subsequent formation and growth of CaMKIIα clusters. We aim to overcome this limitation by designing a more suitable HS-AFM substrate that minimizes non-specific adsorption. Improved substrates and imaging conditions should enable direct visualization of the reversible cluster dynamics expected to occur in neurons.

In conclusion, single-molecule imaging at the mesoscopic scale using HS-AFM directly visualized CaMKIIα clusters formed by CaMKIIα-CaMKIIα intermolecular interactions. Formation of worm-like chain clusters has the potential to be essential for the normal neuronal morphology and proper LTP induction. These data advance our understanding of the mechanisms governing synaptic targeting and are expected to inform models of learning and memory and to aid in elucidating the causes of NDDs.

## Methods

### DNA plasmids

Genes encoding rat (*Rattus norvegicus*) CaMKIIα, CaMKIIα_T286A_, CaMKIIα_T305A/T306V_ and calmodulin (CaM), were prepared as described previously^37^. Point mutations corresponding to CaMKIIα_P212L_ and CaMKIIα_E183V_ were introduced using either a QuikChange site-directed mutagenesis kit (Agilent Technologies, USA) or KOD One^TM^ PCR Master Mix (Blue) (TOYOBO, Japan). For all CaMKIIα constructs, 6×His/Strep tags (MDYKDDDD<u>HHHHHH</u>K<u>WSHPQFEK</u>GTGGQQMGRDLYDDDDKDLYKSGLRSRA) were attached to the N-termini and inserted into a modified pEGFP-C1 vector in place of EGFP (Clontech). CaM constructs tagged with 6×His were cloned into the pRSET bacterial expression vector (Invitrogen, USA), with a 6×His tag (MRGS<u>HHHHHH</u>GMASMTGGQQMGRDLYDDDDKDRSEFG) fused to the N-termini.

### Purification of CaM from bacteria

His-tagged CaM was expressed and purified from *Escherichia coli* (DH5α) following the protocol previously described^37^. In brief, the bacterial pellet was dissolved in PBS containing 1% Triton X-100 and 5 mM imidazole, sonicated, and then centrifuged. The resulting supernatant was applied to a Ni^2+^-nitrilotriacetate (NTA) affinity column (HiTrap, GE Healthcare, USA). The bound protein was washed sequentially with buffers containing 5 mM and 50 mM imidazole in 20 mM Tris (pH7.4), 150 mM KCl, and eluted with a buffer containing 20 mM Tris-HCl (pH7.4) and 500 mM imidazole. Protein concentration was determined using the Bradford assay (Bio-Rad, USA) with BSA as a standard. The resulting protein concentration was typically approximately 500 µM in a volume of 1000 μl. The purified CaM was stored at 4°C or −30°C in a buffer containing 20 mM Tris-HCl (pH7.4), 150 mM KCl, and 5 mM DTT.

### Purification of CaMKIIα from HEK293 cells

All CaMKIIα proteins were purified from HEK293 or HEK293T cells following the protocol previously described^37^. Briefly, 6×His/Strep-tagged CaMKIIα proteins were expressed by transfecting cells with plasmids encoding the target genes using Lipofectamine 3000 (Thermo Fisher Scientific, USA) according to the manufacturer’s instructions. After 24 hours, the cells were lysed in a cold buffer containing 1% Triton X-100, 5% glycerol, 50 mM Tris, 150 mM KCl, and 4 mM EDTA (pH adjusted to 7.6) and then centrifuged at 20,000 × g for 10 min. The supernatant was loaded onto Strep-Tactin Sepharose (Nacalai, Japan), washed with 20 mM Tris-HCl, 150 mM KCl, and eluted with 2.5 mM desthiobiotin or D-biotin in the same buffer. Further purification was performed using an NTA column. Protein concentrations were determined using the Bradford assay (Bio-Rad, USA), with BSA as a standard. Purity was verified by silver staining according to the manufacturer’s protocol (COSMO BIO, Japan). Typically, final concentrations ranged from 0.2 µM to 10 µM in a total volume of 200–500 μL. The purified proteins were stored at −30°C in 20 mM Tris-HCl (pH 7.4), 150 mM KCl, 40% glycerol, and 5 mM DTT.

### Biochemical assay

The kinase assay was performed using purified proteins at concentrations of 300 nM CaMKIIα_WT_, CaMKIIα_T286A_, CaMKIIα_T305A/T306V_, and 100 nM CaMKIIα_P212L_. For the Ca^2+^/CaM-bound state, CaMKIIα proteins were mixed with CaM at a 1:1 molar ratio in a reaction buffer containing 50 mM Tris-HCl (pH 7.4), 150 mM KCl, 10 mM MgCl_2_, and 1 mM CaCl_2_. For phosphorylation at T286, CaMKIIα was similarly mixed with CaM at a 1:1 molar ratio in the same reaction buffer, supplemented with 1 mM ATP. For both conditions, reaction mixtures were incubated for 5 min at 30°C. To dissociate Ca^2+^/CaM from CaMKIIα, 2 mM EGTA was added, followed by a 5 min incubation at 30°C. Subsequently, the samples were incubated for an additional 10 min at 25°C to align with the conditions used for HS-AFM observations. The reactions were terminated by adding SDS sample buffer. Western blotting was conducted using the following primary antibodies: anti-phospho-CaMKII (Thr286) (D21E4; Cell Signaling Technology, USA), anti-CaMKIIα Phospho-Thr305 (abx012403; Abbexa), anti-phospho-CaMKII (Thr306) (p1005-306; PhosphoSolutions, USA), anti-His-tag (27E8; Cell Signaling Technology, USA), and IRDye 800CW conjugated anti-rabbit/mouse secondary antibodies (LI-COR, USA).

### HS-AFM equipment

HS-AFM observations were performed using a custom-built instrument^37,41^. The deflection of the cantilever (AC-7, 8, or 10, MNOIC, Japan) was detected via an optical beam deflection system employing an infrared (IR) laser at 780 nm with a power of 0.8 mW. The IR laser was focused on the back of the cantilever through a 20× objective lens (CFI S Plan Fluor LWD 20X, Nikon, Japan). The cantilever’s spring constant was approximately 0.2 N/m, with a resonant frequency around 400 kHz and a quality factor of 2 in liquid. The original HS-AFM tip had a bird-beak-like triangular shape; to improve the spatial resolution of HS-AFM, an amorphous carbon tip was fabricated on it via electron beam deposition (EBD) using a scanning electron microscope (FE-SEM; Verios 5UC, Thermo Fisher Scientific, USA). The additional EBD tip measured about 250 nm in length, with the apex radius reduced to approximately 1 nm after plasma etching with a plasma cleaner (Tergeo, P.I.C. Scientific, USA). The cantilever’s amplitude was set below 1 nm, with the setpoint maintained at 90% of this amplitude for tip-sample distance control. Image acquisition and tip-scanning were controlled by the custom software UMEX. To minimize sample damage caused by tip scanning, an “only trace imaging” mode^62^ was used, in which images were acquired only during the forward scan, while during the retrace, the tip was lifted slightly off the surface and scanned at the same speed without image acquisition.

### HS-AFM observations

HS-AFM observations of CaMKIIα_WT_, CaMKIIα_T286A_, CaMKIIα_T305A/T306V_, and CaMKIIα_P212L_ were performed on bare mica. To visualize CaMKIIα_E183V_ and to capture magnification images of interacting holoenzymes, experiments used mica modified with cationic C2 pillar[5]arene (P[5]A+), which renders the surface positively charged, as described previously^37^.

[Condition 1]

To observe cluster formation dynamics in the basal state, 600 nM CaMKIIα was added to imaging buffer A (50 mM Tris-HCl (pH 7.4), 150 mM KCl, and 10 mM MgCl_2_) during HS-AFM scanning. For the Ca^2+^/CaM-bound state, CaMKIIα and Ca^2+^/CaM were premixed at a 1:1 molar ratio in buffer A with 1 mM CaCl_2_ and incubated for 5 min at 30°C. Thereafter, 300 nM of the premixed Ca^2+^/CaM-bound CaMKIIα was added to the imaging buffer A with 1 mM CaCl_2_ during HS-AFM scanning.

[Condition 2]

To observe stable CaMKIIα clusters, a 3 μL drop of CaMKIIα was deposited onto a mica surface and incubated for 5 min. The surface was washed five times with 20 μL of the appropriate imaging buffer before HS-AFM observations. Protein concentration was adjusted to yield 30–50 holoenzymes within a 300 × 300 nm^2^ imaging area. Observations of the basal states of CaMKIIα_WT_, CaMKIIα_T286A_, CaMKIIα_T305A/T306V_, and CaMKIIα_P212L_ were performed in imaging buffer A. For Ca^2+^/CaM-bound CaMKIIα, 300 nM CaMKIIα was premixed with CaM (1:1 molar ratio) in imaging buffer A with 1 mM CaCl_2_ and incubated at 30°C for 5 min; HS-AFM observations were performed in imaging buffer A with 1 mM CaCl_2_. For Ca^2+^/CaM-bound CaMKIIα with ATP, 150–300 nM CaMKIIα was premixed with CaM (1:1 molar ratio) in imaging buffer A with 1 mM CaCl_2_ and 1 mM ATP, incubated at 30°C for 5 min; HS-AFM observations were performed in imaging buffer A with 1 mM CaCl_2_ and 1 mM ATP. For the fully phosphorylated state, 200–600 nM CaMKIIα was premixed with CaM (1:1 molar ratio) in imaging buffer A with 1 mM CaCl_2_ and 1 mM ATP, incubated at 30°C for 5 min; 2 mM EGTA was then added to dissociate Ca^2+^/CaM, followed by further 5 min incubation at 30°C. HS‒AFM observations were then performed in imaging buffer A with 1 mM ATP.

[Condition 3]

The observations of CaMKIIα_E183V_ were performed in imaging buffer B (50 mM Tris-HCl (pH 7.4), 15 mM KCl, 10 mM MgCl_2_). Under these experimental conditions (characterized by low KCl concentration), the interaction between CaMKIIα and the AFM substrate is strengthened, facilitating accurate observation of the holoenzyme structure in a relatively fixed state. Similarly, all HS-AFM observations of enlarged contact regions between two CaMKIIα holoenzymes were performed in imaging buffer B containing CaCl_2_ or ATP, depending on the experimental conditions (as above). Both CaMKIIα and CaM were premixed at 50 nM each before imaging.

### HS-AFM image processing and data analysis

HS-AFM images were processed and analyzed using Fiji (ImageJ) software (NIH, USA)^63^. Background subtraction was performed on each image using the “Subtract Background” function with a rolling-ball radius of 150–400 pixels. Lateral drift across sequential HS-AFM images was corrected using the “Template Matching and Slice Alignment” plugin^64^. To extract the coordinates of CaMKIIα holoenzymes in HS-AFM videos, the “MTrackJ” plugin^65^ was used for semi-automated tracking of the intensity-weighted mean location of each CaMKIIα hub assembly. Trajectories of CaMKIIα holoenzymes were analyzed in the basal state (13,107 points across 170 frames from 604 holoenzymes) and in the Ca^2+^/CaM-bound state (16,925 points across 250 frames from 586 holoenzymes). Quantitative analysis of holoenzyme counts within clusters included: for CaMKIIα_WT_, 491 holoenzymes in 10 images (basal), 459 in 11 images (Ca^2+^/CaM), 380 in 10 images (Ca^2+^/CaM+ATP), and 376 in 10 images (EGTA post Ca^2+^/CaM/ATP). For CaMKIIα_T286A_, 465 holoenzymes in 10 images (basal), 446 in 11 images (Ca^2+^/CaM+ATP), and 339 in 7 images (EGTA post Ca^2+^/CaM/ATP). For CaMKIIα_T305A/T306V_, 536 holoenzymes in 12 images (basal), 507 in 11 images (Ca^2+^/CaM+ATP), and 430 in 10 images (EGTA post Ca^2+^/CaM/ATP). For CaMKIIα_P212L_, 656 holoenzymes in 14 images (basal), 329 in 8 images (Ca^2+^/CaM+ATP), and 596 in 11 images (EGTA post Ca^2+^/CaM/ATP).

### Definition of cluster formation

CaMKIIα clustering during HS-AFM observations was analyzed using the Density-Based Spatial Clustering of Applications with Noise (DBSCAN) algorithm^66,67^ applied to the coordinates of individual holoenzymes in each frame. DBSCAN requires two parameters: a distance threshold (eps) and a minimum number of points (min_samples). In this study, eps was set to 13 nm for the basal state and 16 nm for the Ca^2+^/CaM-bound state. These values were calculated from reported *D*_h-k_ (distance between the center of the hub assembly and the kinase domains) of 10.6 nm (basal) and 13.6 nm (Ca^2+^/CaM-bound)^37^, adding the EM^12^-determined kinase-domain radius of 2.3 nm. min-sample was set to 2 to detect clusters comprising two or more adjacent CaMKIIα holoenzymes.

### Mean square displacement (MSD)

To assess CaMKIIα holoenzyme mobility on mica surface in clustered versus nonclustered states, we tracked the hub-assembly center coordinates of Ca^2+^/CaM-bound CaMKIIα_WT_ and calculated MSDs. DBSCAN was applied per frame to classify each holoenzyme as in-cluster or out-of-cluster; cluster duration thresholds of 15, 50, 75, 100, 125, and 150 s were used. Here, holoenzymes that remained in a cluster for longer than the chosen threshold were designated “in-cluster”, whereas those that remained outside of a cluster for longer than the threshold were designated “out-of-cluster”. MSD was computed separately for each group. Although MSD typically provides reliable measures of mobility with extended recording and large particle counts, the transient nature of CaMKIIα clustering in our data limited the number of holoenzymes that remained clustered over long intervals. Consequently, we quantified holoenzyme mobility for in-cluster and out-of-cluster populations across six different cluster-duration thresholds and analyzed the threshold-dependent MSD behavior (Supplementary Fig. 2f). Lateral drift correction was performed by subtracting the average frame-to-frame displacement of all holoenzymes from each trajectory. Data from two independent HS-AFM experiments were analyzed. The MSD was calculated using the following equation^68^:

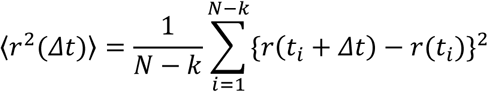

where *r*(*t*_*i*_) represents the two-dimensional position of the holoenzyme at time *t*_*i*_, *k* is an integer, Δ*t* is the time lag defined as *k* × *δt*, where *δt* is the time interval between frames, and *N* is the total number of frames in the trajectory. MSDs were fitted using the following equation^69^:

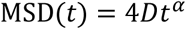

where *D* is the diffusion coefficient and *⍺* is the anomalous diffusion exponent. The fitting range was set from 0 s to half of the corresponding duration threshold.

### Radial distribution function (RDF)

To quantify the spatial distribution of CaMKIIα holoenzymes on the AFM substrate, pairwise distances between all holoenzymes within each frame were calculated from their coordinates, and the radial distribution function *g*(*r*) was computed using the following equation^50^.

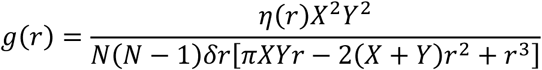

where *r* is the distance between holoenzymes, *η*(*r*) is the number of holoenzyme pairs separated by a distance *r* ± *δr*/2 (*δr* = 2 nm), *N* is the total number of holoenzymes within the frame, and *X* and *Y* are spatial dimensions of the observation area (300 × 300 nm^2^). The term in brackets in the denominator accounts for boundary effects due to the finite observation area. To derive an averaged radial distribution function across frames, the *g*(*r*) values were combined using a weighted average:

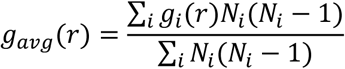

The summation runs over all frames *i*, with *N*_*i*_ representing the number of holoenzymes in frame *i*. The average *g*_*avg*_(*r*) was converted to a free energy *w*(*r*) according to the equation:

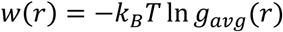

where *k*_*B*_ is the Boltzmann constant and *T* is the absolute temperature, set to 305 K. To correct for distance errors due to variations in AFM tip, the major axis distance of the randomly selected five hub assemblies were normalized based on the 11.0 nm diameter from EM^12^. Coordinates of CaMKIIα_WT_ holoenzymes were used to compute RDFs for the basal state (2,275 points in 54 frames), the Ca^2+^/CaM-bound state (2,524 points in 60 frames), the Ca^2+^/CaM-bound+ATP state (2,008 points in 52 frames), and the fully phosphorylated state (2,362 points in 60 frames). The robustness of the RDF and free-energy profile was assessed by recalculating the RDF using half of the HS-AFM data (Supplementary Fig. 3c, d).

Furthermore, we assessed the stability of CaMKIIα clusters by calculating the free energy per cluster under different activation states. To estimate the free energy of a cluster (*E*_cluster_), all pairwise inter-holoenzyme distances within each cluster (*r_ij_*, where *i* and *j* stand for holoenzyme *i* and *j* in the cluster) were measured and summed pairwise free energy 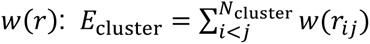, where *N*_cluster_ is the number of holoenzymes in the cluster. The *E*_cluster_ was then divided by *N*_cluster_ to yield the free energy per holoenzyme (*E*_oligomer_): *E*_oligomer_ = *E*_cluster_ / *N*_cluster_ (Supplementary Fig. 3b).

### Simulation of CaMKIIα cluster dynamics in a two-dimensional space

To investigate the factors contributing to the growth of CaMKIIα clusters, particle simulations were conducted on a two-dimensional plane in a 500 × 500 nm^2^ area. Python code for the simulation was developed in-house. Here, a particle corresponds to a CaMKIIα holoenzyme. The particle radii were estimated based on the distance between the hub and the kinase domains observed by single-molecule HS-AFM of CaMKIIα^37^ and the diameter of the kinase domain observed by electron microscopy^12^. The radius for the basal state was set to 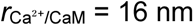.

The binding rate constant of particles to the substrate (*k*_on_) and the dissociation rate constant from the substrate (*k*_off_) were estimated based on HS-AFM observations. Three independent HS-AFM datasets were analyzed. The number of particles on the substrate against observation time was fitted by the exponential function, *A*{1 − *exp*(−*kt*)}, where *A* is the number of particles in the steady state, *k* is the rate constant, *t* is time. During HS-AFM measurements, a slight delay can occur between the onset of particle adsorption onto the substrate and the start of image acquisition. To account for this offset and align the time origins among datasets, the fitting was performed using a corrected time variable *t* + *C*_i_, where *C*_1_ = 14 s, *C*_2_ = 60 s, and *C*_3_ = 0 s for the respective datasets, where *C*_i_ was chosen to minimize the difference from the exponential function. Fitting the combined datasets yielded *A* of 97.8 and *k* of 0.0064 s^−1^ (Supplementary Fig. 6). The dissociation rate constant *k*_off_ was then determined from the experimentally measured dissociation frequency during the steady state, defined as 300 s after data acquisition began. The frequency was the number of particles that detached from the substrate and disappeared from the observation field divided by the observation time. Given a dissociation frequency of 0.1341 s^−1^ and assuming that the number of particles present on the substrate at steady state is *A*, the dissociation rate constant was calculated as *k*_off_ = 0.1341 / *A*, yielding *k*_off_ = 0.0014 s^−1^. Because the rate constant *k* obtained from fitting in a two-state model is the sum of the binding and dissociation rates (*k* = *k*_on_ + *k*_off_), *k*_on_ was calculated to be 0.0050 s^−1^.

Using the obtained values of *A*, *k*_on_, and *k*_off_, the total number of particles within the system, *N*_total_, was estimated. At equilibrium state, the rate of particle binding to the substrate is balanced by the rate of dissociation. Denoting the number of particles freely diffusing in solution as *N*_off_, this balance is described by *N*_off_ × *k*_on_ = *A* × *k*_off_. Because *N*_total_ = *N*_off_ + *A*, the total particle number can be expressed as *N*_total_ = *A* (*k*_on_ + *k*_off_) / *k*_on_. Substituting the experimentally determined parameters into this relation yielded *N*_total_ ≈ 125.1. This value was subsequently adjusted to 135 so that the simulation results more closely reproduced the HS-AFM observations. To validate the chosen parameters, time-dependent curves obtained from 20 independent simulation trials were also fitted with *A*{1 − *exp*(−*kt*)}, resulting in *A* = 93.8 and *k* = 0.0065 s^−1^. The close agreements with the experimental values indicate that the chosen values of *k*_on_, *k*_off_, and *N*_total_ are appropriate (Supplementary Fig. 6).

The particles were allowed to randomly walk. The simulation time step (Δ*t*) was set to 1 s. The particle displacement in the simulation step was adjusted so that the diffusion coefficient (*D*) obtained from the MSD analysis matched the *D* measured in HS-AFM experiments. The MSD in the simulation was estimated using over 12,000 particles. Linear fitting using MSD = 4*Dt* was performed for particles observed for more than 200 s, with the fitted time range was 0–100 s. It was found that *D* in the simulation was close to that in the experiments when the particle displacement followed a normal distribution centered at 1.8 nm/step with a standard deviation of 1 nm (Supplementary Fig. 6). The maximum displacement was set at 10 nm. The direction of movement was randomly assigned based on a uniform distribution. When particles moved outside the simulation area, they were removed from the simulation (open boundary condition). When two particles approached within 32 nm and unnaturally overlapped after particle displacement, both particles were slightly displaced in opposite directions. This adjustment was iteratively performed until all overlaps were eliminated.

Clusters are defined as formed when two particles are within the particle diameter after displacement. To avoid numerical artifacts arising from rounding errors in floating-point calculations, this distance criterion was evaluated with a small offset of 10^−12^ nm added to the threshold. After cluster formation, particles belonging to the same cluster were assumed to move rigidly, such that all particles in a given cluster shared the same displacement magnitude and direction, which were determined by the vector sum of the individual displacement vectors of the particles within the cluster. To estimate the dissociation rate of CaMKII from clusters, the duration during which CaMKII molecules remained in clusters was measured at low density (approximately 60% of the maximum particle number) in HS-AFM observations. The histogram of the duration was fitted with an exponential function, *B*exp(−*λt*), where *λ* is the dissociation rate and was 0.1416 s^−1^.

The dissociation probability per one simulation step (*P*) was derived from *λ* obtained from HS-AFM measurements under low-density conditions, where clusters predominantly exist as dimers. Let *P* denote the dissociation probability of an individual particle from a cluster, and let *P’* denote the probability that a cluster dissociates. Assuming dissociation follows a Poisson process, *P’* was calculated using the relation *P’* = 1 − exp(−*λ*Δ*t*). A dimeric cluster dissociates when either of the two constituent particles dissociates. If dissociation from clusters is independent, the probability that the cluster remains is given by (1 − *P*)^2^, which is the probability that neither particle dissociates. Accordingly, the probability that a cluster dissociates within 1 s observed in HS-AFM observations, denoted as *P′*, is calculated by *P′* = 1 − (1 − *P*)^2^ = 2*P* – *P*^2^. By substituting the experimentally determined *P*′ into this relation, the per-particle dissociation probability *P* was estimated (*P* = 0.0683), which corresponds to the dissociation rate (0.0708 s^−1^) that should be used in the simulation. However, an adjustment to this rate was required to match the simulated dissociation rate from clusters with that observed in HS-AFM experiments, yielding a corrected rate of 0.1054 s⁻¹ (*P* = 0.10).

In simulations including immobile particles, 13 particles were fixed at the coordinates (0, 0), (62.5, 62.5), (−62.5, 62.5), (62.5, −62.5), (−62.5, −62.5), (125, 125), (125, 0), (125, −125), (0, −125), (−125, −125), (−125, 0), (−125, 125), and (0, 125) throughout the simulation. When mobile particles formed a rigid cluster with a fixed particle, the cluster’s motion became rotational, since the fixed particle prevents translation. To account for this rotation, the center of mass of the mobile particles within the cluster was first calculated, and the distance between this center of mass and the fixed particle was defined as the rotation radius. A displacement distance chosen from a normal distribution centered at 1.8 nm per step with a standard deviation of 1 nm was converted into an angular increment along the circumference to ensure that the center of mass displacement follows the normal distribution. The direction of rotation, clockwise or counterclockwise, was chosen randomly. The positions of all mobile particles in the cluster were rotated around the fixed particle.

### Quantification and statistical analysis

Statistical analyses were conducted using Igor Pro 10 (WaveMetrics, USA) or OriginPro2025 (OriginLab, USA). A significance threshold of α = 0.05 was maintained for all tests, with *P* values corrected for multiple comparisons when applicable. Normality of data was tested with the Shapiro‒Wilk test unless otherwise specified. Homogeneity of variances among groups was checked using Levene’s test. When normality was not confirmed, the Kruskal-Wallis test was used for group comparisons. Pairwise comparisons between two groups were performed using the Brunner-Munzel test and Welch’s t-test. For comparisons involving multiple conditions, one-way ANOVA and the Kruskal-Wallis test were used to analyze multiple groups with a single independent variable. As a follow-up test to the Kruskal-Wallis test, Dunn’s test was used to compare every mean with every other mean. The figure captions specify the statistical tests used. *P* > 0.05, not significant (NS); \**P* < 0.05, \*\**P* < 0.01, and \*\*\**P* < 0.001 were considered to indicate statistical significance. The data are presented as the means ± SDs.

## Use of large language models

ChatGPT was employed for proofreading the text.

## Data availability

The data that support the findings of this study are available from the corresponding author upon request.

## Acknowledgments

We thank R. Yasuda (Max Planck Florida Institute) and T. Ando (Kanazawa University) for providing the HS-AFM apparatus; Y. Mikami and Y. Okada (Kanazawa University) for collecting and analyzing the HS-AFM data; Y. Kamide (Kanazawa University) for preparing the CaMKIIα samples. This research was supported by the World Premier International Research Center Initiative (WPI), MEXT, Japan (to M.S.), JSPS KAKENHI grant numbers JP24K21942 (to M.S.) and JP25H00972 (to M.S.), JP22H04926 Advanced Bioimaging Support (ABiS) (to M.S.), 23H0424 (to H.M.), 24H01298 (to H.M.), the Mochida Memorial Foundation for Medical and Pharmaceutical Research (to M.S.), the Uehara Memorial Foundation (to M.S.), the Naito Foundation (to M.S.), JST CREST (JPMJCR1762 to N.K.), JST SPRING (JPMJSP2135 to T.S. and K.M.), and JST ERATO (JPMJER2403 to M.S.).

## Author contributions

Conceptualization: M.S.; methodology: T.S., T.Sumikama, K.M., K.H., U.K., N.K., and M.S.; software: T.S., K.M., and M.S.; validation: T.S., T.Sumikama, K.M., K.H., A.S., H.M., T.D., and M.S.; formal analysis: T.S., T. Sumikama, K.M., T.D., C.B., and M.S.; investigation: T.S., K.M., K.H., H.M., and M.S.; resources: H.M., and M.S.; data curation: M.S.; writing – original draft: M.S.; writing – review & editing: all authors; visualization: T.S., K.M., and M.S.; supervision: T. Sumikama, A.S., C.B., H.M., and M.S.; project administration: M.S.; funding acquisition: H.M., and M.S.

