## Supplemental Figs for "CaMKIIα holoenzymes self-organize into worm-like mesoscale clusters"

Includes: Supplementary Figures 1–8, Supplementary Discussion, and Supplementary Movies 1–12.

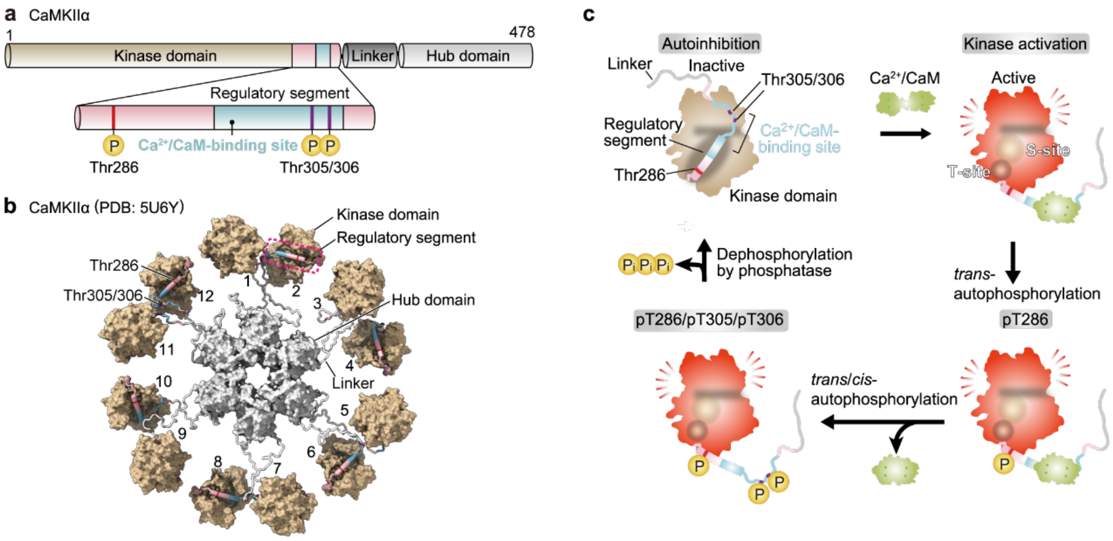


**Supplementary Fig. 1: Illustration of CaMKIIα structure.**

**a,** Domain structure of CaMKIIα, with numbers indicating amino acid sequence positions. **b,** Pseudoatomic model of the CaMKIIα dodecamer derived from EM data[12]. The molecular surface is depicted as a solvent-excluded surface, highlighting individual structural elements: the hub domain (light gray, residues 345–472), kinase domain (tan, residues 1–273), regulatory segment (magenta cylinder, residues 274–314) including the $Ca^{2+}$/CaM-binding region (blue cylinder, residues 293–310), and the linker region (white ribbon, residues 315–344). Phosphorylation sites at T286, T305, and T306 are marked in red and purple.

**c,** Conformational changes during CaMKIIα activation, highlighting a single kinase domain. The kinase domain is autoinhibited by its regulatory segment in the inactive state. The binding of $Ca^{2+}$/CaM releases the regulatory segment, revealing the substrate-binding site (S-site) and the GluN2B-binding site (T-site), resulting in kinase activation. Activation of neighboring kinase domains causes autophosphorylation at T286 (pT286). Further autophosphorylation at T305/T306 (pT286/pT305/pT306) occurs after $Ca^{2+}$/CaM dissociation. Dephosphorylation by protein phosphatases returns the kinase domain to its inactive state.

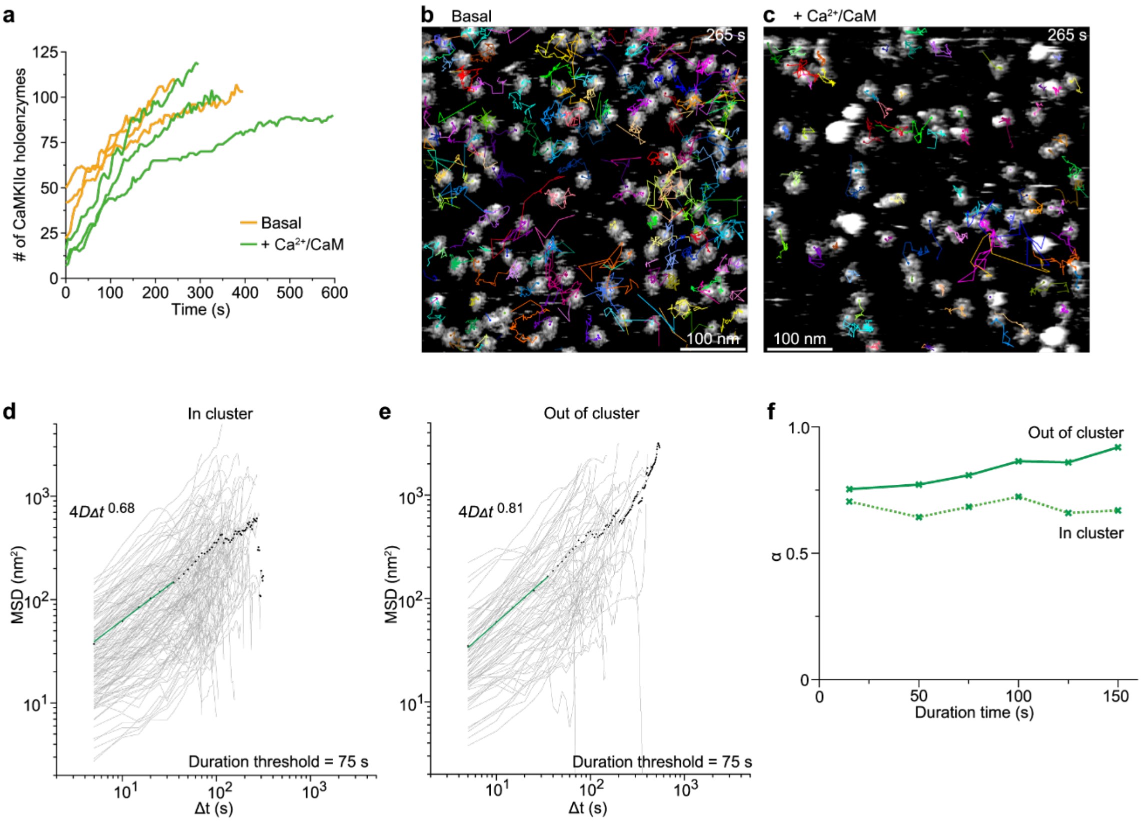


**Supplementary Fig. 2: Restricted mobility of CaMKIIα holoenzymes within clusters.**

**a,** Time-course of the number of CaMKIIα holoenzymes appearing within a 500 × 500 $nm^2$ area under the basal and $Ca^{2+}$/CaM-bound states. Data from three independent experiments are plotted as separate lines.

**b, c,** Representative trajectories of hub assembly coordinates of CaMKIIα holoenzymes in the basal (**b;** Supplementary Video 3) and $Ca^{2+}$/CaM-bound (**c;** Supplementary Video 4) states, tracked over 265 s.

**d,e**, MSD of the $Ca^{2+}$/CaM-bound CaMKIIα holoenzymes. MSDs were calculated from trajectories classified as “in cluster” (**d**) and “freely diffusing” (Out of cluster, **e**) across three independent experiments. Gray lines illustrate individual MSD; black dots mark the mean MSD values. Fitting curves are shown in green.

**f**, The relationship between the exponent α and the duration time for cluster classification (see Methods). Increasing the duration threshold time for freely diffusing holoenzymes (out of cluster) led α to approach 1. Conversely, for holoenzymes within clusters (in cluster), α remained relatively constant, confirming constrained mobility in clusters.

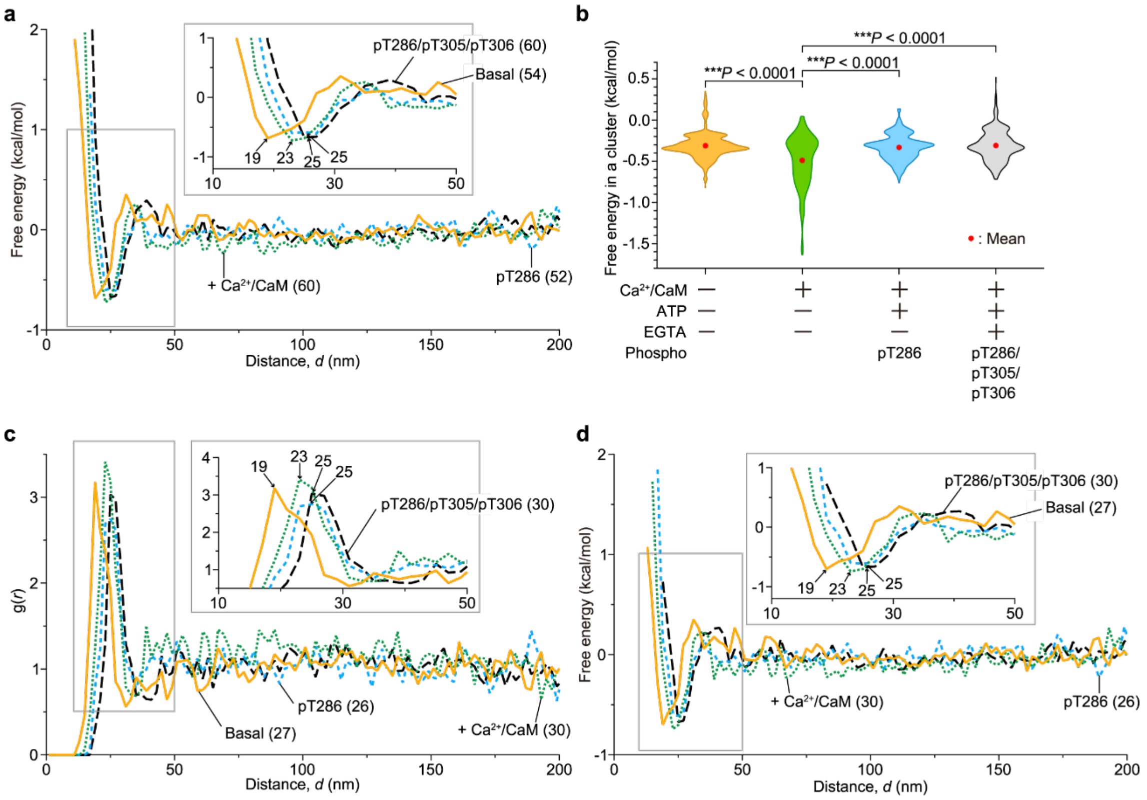


**Supplementary Fig. 3: Free energy of CaMKIIα clusters.**

**a,** Free-energy profile of CaMKIIα holoenzymes converted from the RDF shown in Fig. 3e (see Methods). The inset at upper right shows a magnified view of the region bounded by the gray box and parentheses indicate the number of images used for analysis (also in **c**,**d**). Bin width, 2 nm.

**b,** Calculated free energy per holoenzyme within a cluster under different experimental conditions. Free energy of holoenzyme pairs was computed from the profiles in (**a**) (see Methods). Numbers of analyzed clusters: $n$ = 555 (basal state), 447 ($Ca^{2+}$/CaM-bound), 422 (pT286), and 471 (pT286/pT305/pT306). Kruskal–Wallis test with Dunn's post hoc test. All tests are two-tailed. NS, not significant ($P > 0.05$); ***$P < 0.001$.

**c,d,** Assessment of RDF and free-energy profile robustness. RDFs (**c**) and the resulting free-energy profiles (**d**) were recomputed using the first half of sequential HS-AFM frames from each experiment. The profiles from these truncated datasets closely match those from the full datasets, supporting the robustness of the original analyses.

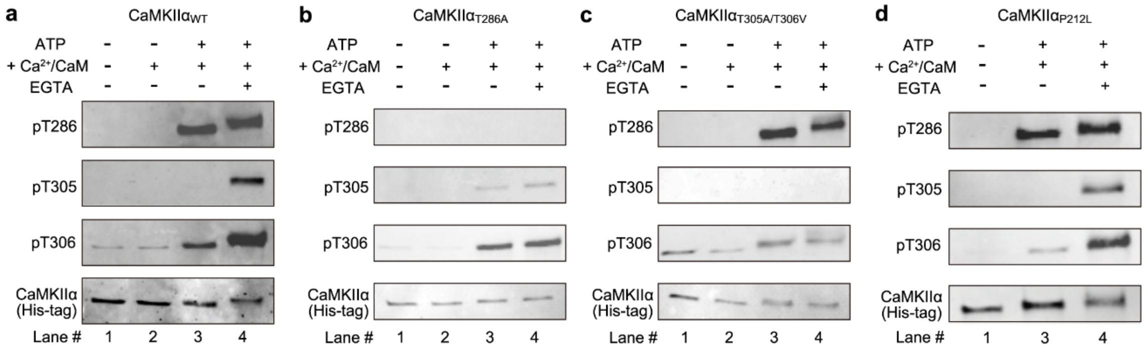


**Supplementary Fig. 4: Western blots of phosphorylation states of purified CaMKIIα.**
**a–d,** (Lane #1) Purified proteins were loaded without activation.
(Lane #2 in **a–c**) CaMKIIα proteins (300 nM) incubated with 300 nM CaM in reaction buffer (50 mM Tris-HCl (pH 7.4), 150 mM KCl, and 10 mM $MgCl_2$) with 1 mM $CaCl_2$ at 30°C for 5 min, showing no phosphorylation.
(Lane #3 in **a–d**) CaMKIIα proteins (300 nM, or 100 nM for P212L) were incubated with CaM (300 nM, or 100 nM for P212L) in reaction buffer containing 1 mM $CaCl_2$ and 1 mM ATP at 30°C for 5 min. This protocol induced phosphorylation at T286.
(Lanes #4 in **a–d**) $Ca^{2+}$/CaM dissociation by incubation with 2 mM EGTA for 5 min at 30°C followed by 10 min at 25 °C. This protocol induced phosphorylation at T286/T305/T306.
The amount of protein loaded was assessed with an anti-His tag antibody.

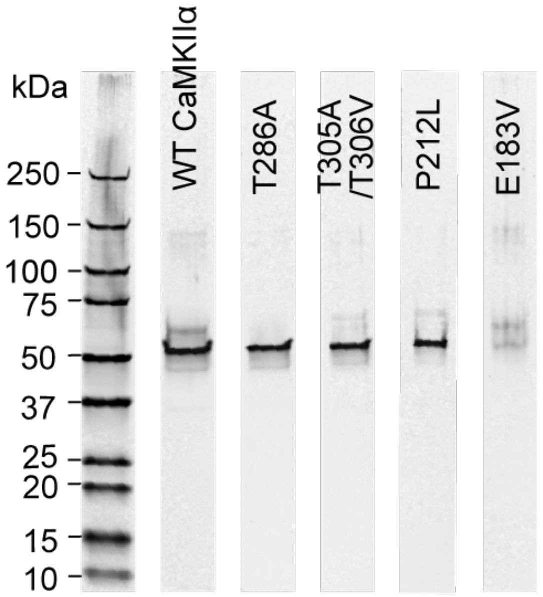


**Supplementary Fig. 5: Silver staining of purified CaMKIIα proteins.**

CaMKII holoenzymes were purified from HEK-293 or HEK-293T cells in the absence of ADP/ATP using a two-step affinity purification via His Strep tags.

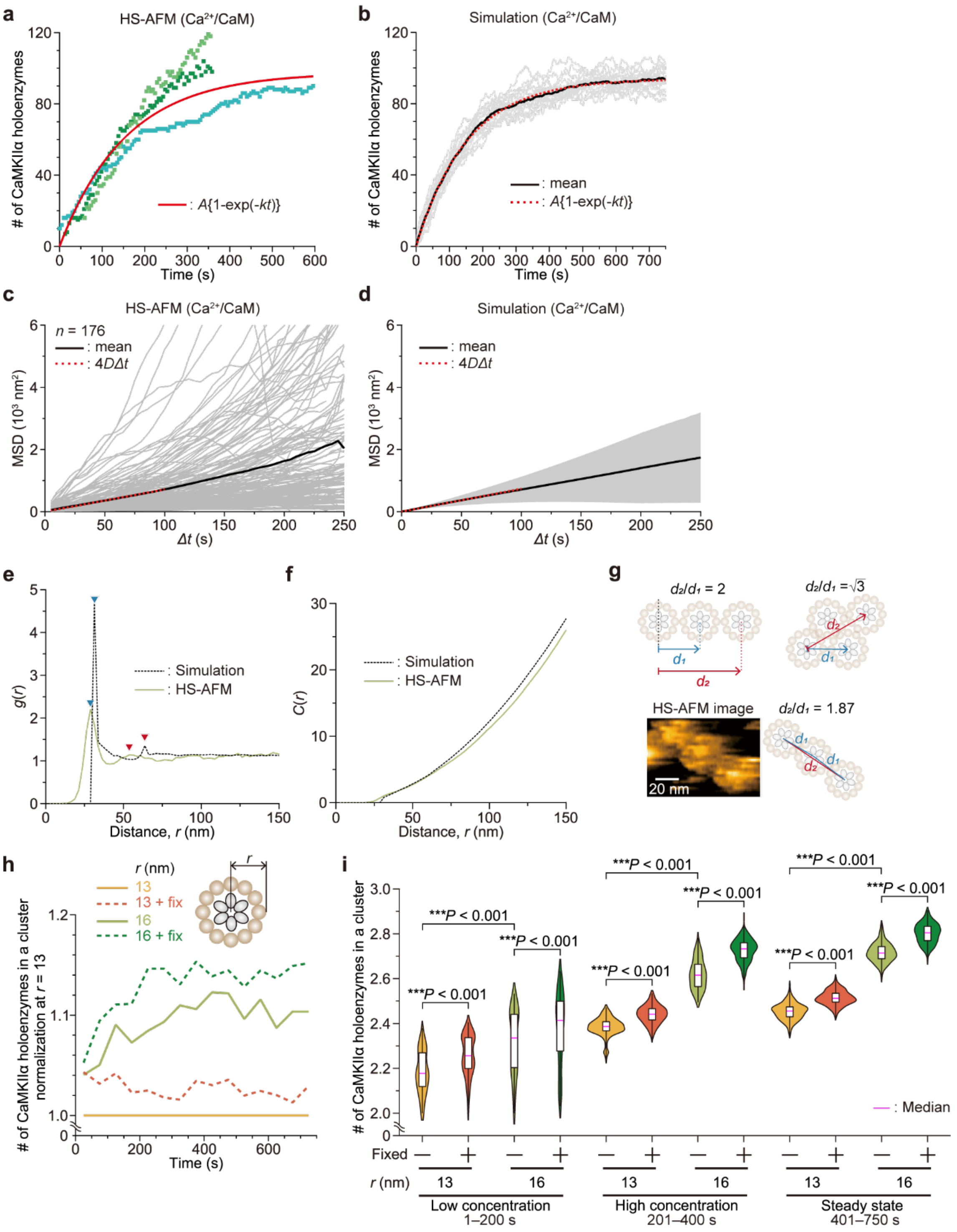


**Supplementary Fig. 6: Simulation of CaMKIIα clustering dynamics in 2D space.**
**a,** Experimental time courses of CaMKIIα holoenzyme numbers measured by HS-AFM. Each colored point is an independent replicate; red solid lines are fit curves. The fitting results are listed in Supplementary Table 2 (also in **b–d**).

**b,** Time course of CaMKIIα holoenzyme numbers in simulations. Individual simulation traces are shown in gray ($n$ = 20), the mean trace in black, and the fit to the mean in red dashed lines.

**c,** MSDs of CaMKIIα holoenzymes observed by HS-AFM. The analysis includes holoenzymes observed continuously for more than 200 s (also in **d**). Each gray line corresponds to an individual holoenzyme. The black line shows the mean MSD; the red dashed line indicates the fit of the mean MSD (also in **d**).

**d,** MSDs of CaMKIIα holoenzymes across 50 independent simulations. The gray shaded regions illustrate the variation (standard deviation).

**e,** RDFs measured in the $Ca^{2+}$/CaM-bound states. Solid green lines indicate experimental data from HS-AFM, while dashed black lines show the results from simulations. The first and second peaks are marked by blue and red arrowheads, respectively. The density of CaMKIIα holoenzymes analyzed varied from 56 to 119 per 500 × 500 $nm^2$ in the $Ca^{2+}$/CaM-bound states. A bin width of 2.5 nm was used.

**f,** Coordination number $C(r)$ computed from RDFs for the $Ca^{2+}$/CaM-bound states. Solid green lines represent data from HS-AFM, while dashed lines indicate simulation results.

**g,** Schematic illustration showing the ratio of the second peak position to the first in the RDF. The ratio is calculated as $d_2/d_1$, where $d_1$ is the distance to the first peak, and $d_2$ is that to the second peak. A ratio of 2.0 suggests a linear arrangement of holoenzymes in clusters, while a ratio of $\sqrt{3}$ implies a two-dimensional close-packed arrangement. (Bottom) Representative HS-AFM images of holoenzymes forming clusters, alongside their corresponding schematic illustration.

**h,** Number of CaMKIIα holoenzymes per cluster normalized at the $r$ = 13 nm. Simulation results over 750 s were segmented into 50 s intervals, and the average number within each interval was plotted. Each condition was independently simulated in 50 repetitions.

**i,** Violin plots with box plots show the distribution of CaMKIIα holoenzymes per cluster, categorized into three temporal phases. The box limits mark the $25^{th}$ and $75^{th}$ percentiles (IQR), and whiskers extend to the minimum and maximum values. Welch's t-test was used to compare. All tests are two-tailed. NS, not significant ($P > 0.05$); ***$P < 0.001$.

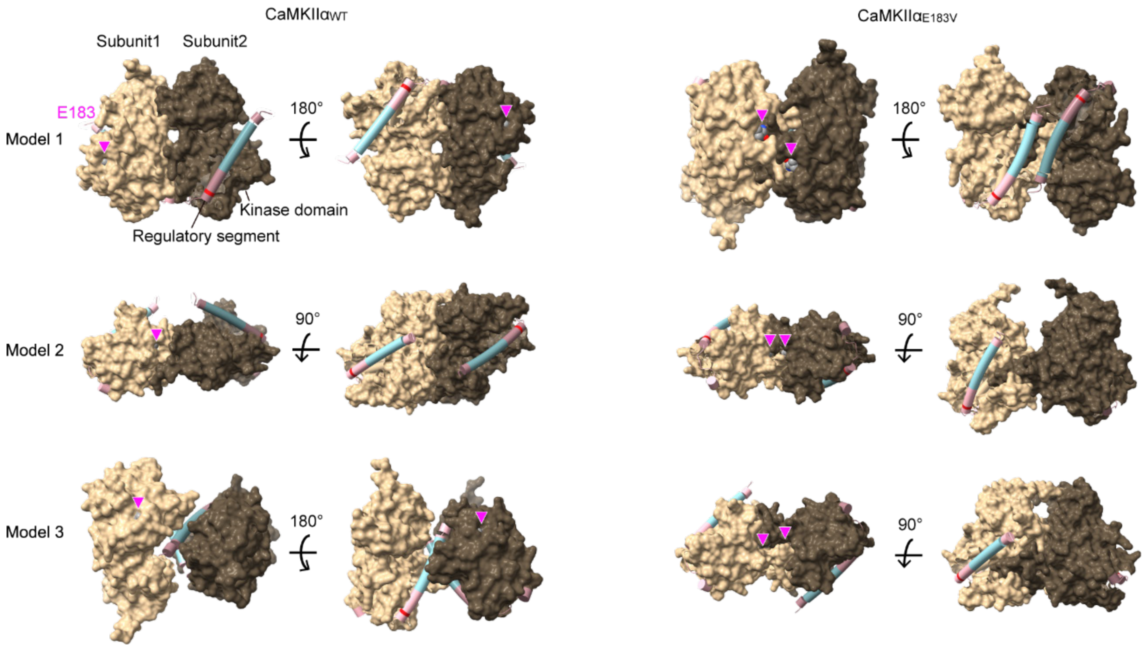


**Supplementary Fig. 7: AF3 prediction for CaMKIIα$_{E183V}$ subunit-subunit interfaces.** Predicted models of intermolecular interactions among CaMKIIα subunits were generated using residues 1–314 (kinase domain with regulatory segment) with AF3 (see also Table 2). Residues 183 is highlighted by magenta arrowheads and depicted as spheres.

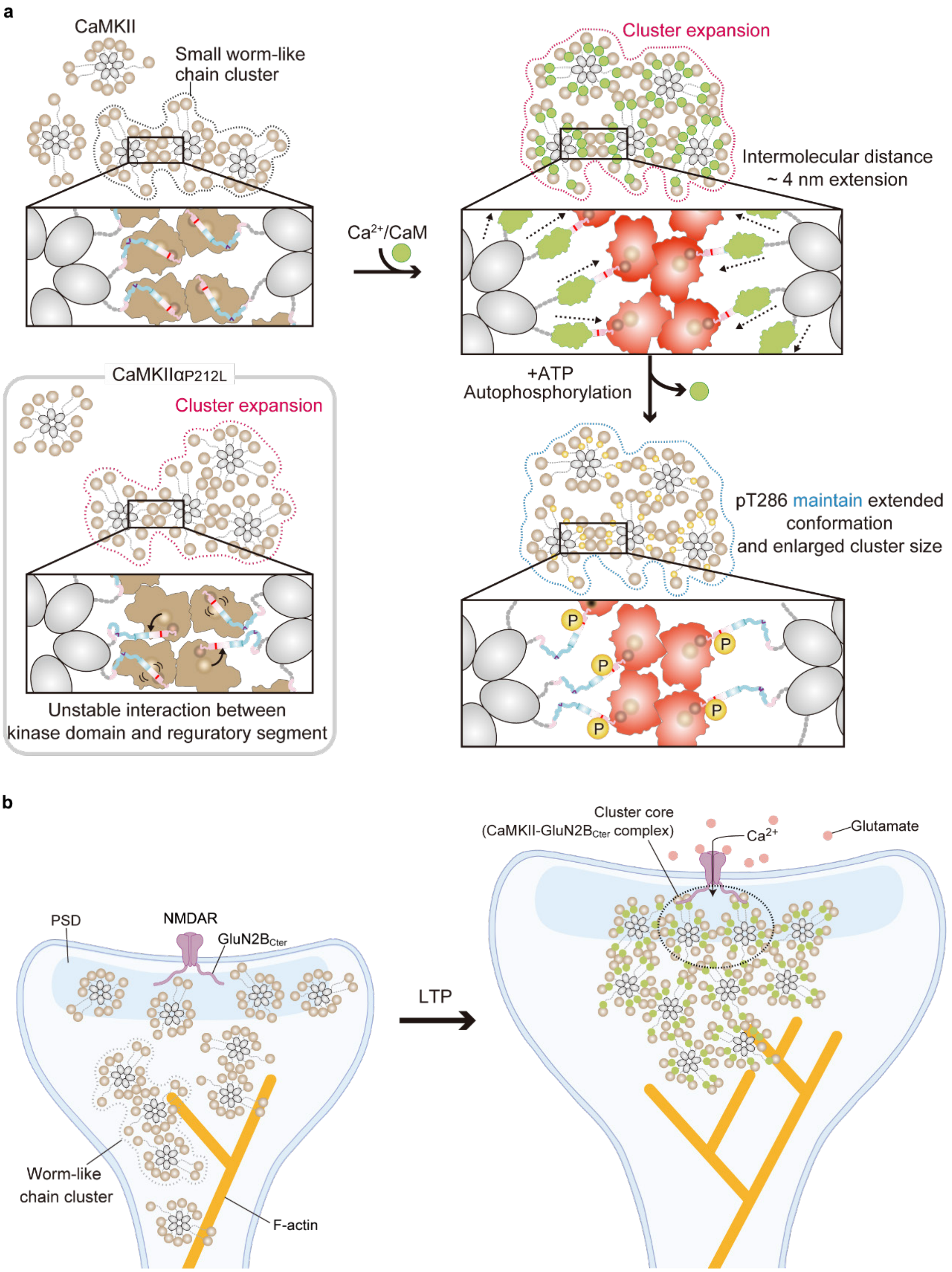


**Supplementary Fig. 8: Proposed model of CaMKII localization in the PSD.**

**a,** Models of CaMKII cluster in the basal state (WT and P212L, left), the $Ca^{2+}$/CaM-bound state (top right), and the pT286 state (bottom right).

**b,** Model of CaMKII distribution within dendritic spines before (left) and after $Ca^{2+}$ stimulation (right).

**Supplementary Discussion**

We computed the MSD and RDF of simulated CaMKIIα holoenzyme trajectories and compared them with experimental measurements to evaluate the accuracy of our simulation in reproducing HS-AFM data. The MSD derived from the simulation, using the parameters in Supplementary Table 1, was in good agreement with the HS-AFM data in the $Ca^{2+}$/CaM-bound state (Supplementary Fig. 6c,d, Supplementary Table 2).

The appearance of RDF calculated using HS-AFM data obtained under the conditions shown in Fig. 2 differed from the simulation results, owing to use of a rigid cluster model in the simulation (Supplementary Fig. 6e). The peaks in RDF from the simulations are narrower and more sharply defined due to the fixed positions of holoenzymes in clusters, whereas in the HS-AFM observations, the peaks are broader due to positional fluctuations of holoenzymes in clusters. To demonstrate that these RDFs are intrinsically similar despite their differing appearances, the coordination number $C(r)$, was computed by integrating RDF (Supplementary Fig. 6f). Within the first peak region (40–45 nm), $C(r)$ was between 1.84–2.29 (in HS-AFM) and 1.85–2.40 (in simulations). Although the difference gradually increased at longer distances around 150 nm, the difference between the two datasets remained below 1.7, supporting their intrinsic similarity.

Furthermore, the ratio of the distances ($d_2/d_1$), where $d_1$ is the distance of the first peak (HS-AFM: 28.75 nm; simulation: 31.25 nm), and $d_2$ is the distance of the second peak (HS-AFM: 53.75 nm; simulation: 63.75 nm), was calculated. This ratio offers insight into the geometric arrangement of holoenzymes in clusters (Supplementary Fig. 6g). In a linear configuration, next-nearest neighbor holoenzymes are positioned approximately twice the distance of the nearest neighbors, resulting in a ratio close to 2.0. If the holoenzymes are closed-packed, the ratio should be $\sqrt{3}$. The measured ratios were 1.87 in HS-AFM data and 2.04 in the simulations. Because the HS-AFM ratio exceeds $\sqrt{3}$. This suggests that most CaMKIIα holoenzymes are likely aligned in a linear arrangement within the clusters, as observed in both experiments and simulations. Consistent with this, linearly arranged holoenzymes are visible in both the HS-AFM data (Supplementary Video 4) and the simulation results (Supplementary Video 10). These findings confirm that the simulations effectively reproduce the HS-AFM observations, including the behaviors of MSD, the $C(r)$, and overall geometric arrangement.

**Supplementary Table 1. Simulation parameters**

| Parameters | |
|---|---|
| Particle radius (nm) | 13 (Basal) or 16 ($Ca^{2+}$/CaM) |
| Step size (nm) | 1.8 |
| $k_{on}$ ($s^{-1}$) | 0.0050 |
| $k_{off}$ ($s^{-1}$) | 0.0014 |
| $N_{total}$ | 135 |
| Dissociation rate ($s^{-1}$) | 0.1054 |

**Supplementary Table 2. Fitting results**

| Parameters | $Ca^{2+}$/CaM-bound state | |
|---|---|---|
| | HS-AFM | Simulation |
| $A$ | 97.8 | 93.8 |
| $k$ ($s^{-1}$) | 0.0064 | 0.0065 |
| $D$ ($nm^2/s$) | 1.8 | 1.8 |

**Supplementary Movie legends**

**Supplementary Movie 1: Three representative HS-AFM videos show CaMKIIα absorption onto the mica surface in the basal state.** CaMKIIα (600 nM) was added to the imaging buffer during HS-AFM scanning. White arrows indicate CaMKIIα holoenzymes, and white dotted lines highlight holoenzymes immediately following adsorption onto the mica surface. Image size, 600 × 200 pixels$^2$ (acquired in HS-AFM observations); scan area, 500 × 500 nm$^2$; frame rate, 0.2 fps.

**Supplementary Movie 2: Three representative HS-AFM videos show CaMKIIα absorption onto the mica surface in the $Ca^{2+}$/CaM-bound state.** CaMKIIα (300 nM), premixed with $Ca^{2+}$/CaM (1 mM $CaCl_2$ and 300 nM CaM), was added to the imaging buffer during HS-AFM scanning. White arrows indicate CaMKIIα holoenzymes, and white dotted lines highlight holoenzymes immediately following adsorption onto the mica surface. Image size, 600 × 200 pixels$^2$ (acquired in HS-AFM observations); scan area, 500 × 500 nm$^2$; frame rate, 0.2 fps.

**Supplementary Movie 3: Three representative HS-AFM videos show the basal state of CaMKIIα in medium-to-high density arrangements on the mica surface.** CaMKIIα (600 nM) was added to the imaging buffer during HS-AFM scanning. White dotted lines outline typical CaMKIIα clusters (see Methods). Right: Trajectory of the hub assembly center coordinates. Image size, 600 × 200 pixels$^2$ (acquired in HS-AFM observations); scan area, 500 × 500 nm$^2$; frame rate, 0.20 fps.

**Supplementary Movie 4: Three representative HS-AFM videos show the $Ca^{2+}$/CaM-bound state of CaMKIIα in medium-to-high density arrangements on the mica surface.** CaMKIIα (300 nM), premixed with $Ca^{2+}$/CaM (1 mM $CaCl_2$ and 300 nM CaM), was added to the imaging buffer during HS-AFM scanning. White arrows indicate the formation of a typical CaMKIIα cluster. Right: Trajectory of the hub assembly center coordinates. Image size, 600 × 200 pixels$^2$ (acquired in HS-AFM observations); scan area, 500 × 500 nm$^2$; frame rate, 0.20 fps.

**Supplementary Movie 5: HS-AFM video of stable CaMKIIα clusters in the basal state on a mica surface.** CaMKIIα (300 nM) was deposited onto the AFM substrate, and HS-AFM observation was performed after incubation for 5 min. The white dotted line indicates a representative cluster. Image size, 900 × 300 pixels$^2$; scan area, 300 × 300 nm$^2$; frame rate, 0.33 fps.

**Supplementary Movie 6: HS-AFM video of stable CaMKIIα clusters in the $Ca^{2+}$/CaM-bound state on a mica surface.** CaMKIIα was activated by $Ca^{2+}$/CaM (1 mM $CaCl_2$ and 300 nM CaM). The white dotted line indicates representative clusters. An enlarged view highlighting intermolecular interactions of CaMKIIα is shown on the right. Image size, 900 × 300 pixels$^2$; scan area, 300 × 300 nm$^2$; frame rate, 0.33 fps.

**Supplementary Movie 7: HS-AFM video of stable CaMKIIα clusters phosphorylated at T286 on a mica surface.**

CaMKIIα was activated by $Ca^{2+}$/CaM and ATP (1 mM $CaCl_2$, 300 nM CaM, and 1 mM ATP). The white dotted line indicates representative clusters. An enlarged view highlighting intermolecular interactions of CaMKIIα is shown on the right. Image size, 900 × 300 pixels$^2$; scan area, 300 × 300 nm$^2$; frame rate, 0.33 fps.

**Supplementary Movie 8: HS-AFM video of stable CaMKIIα clusters fully phosphorylated at T286/T305/T306 on a mica surface.**

CaMKIIα was treated with 2 mM EGTA and 1 mM ATP following stimulation (1 mM $CaCl_2$, 300 nM CaM, and 1 mM ATP). The white dotted line indicates representative clusters. An enlarged view highlighting intermolecular interactions of CaMKIIα is shown on the right. Image size, 900 × 300 $pixels^2$; scan area, 300 × 300 $nm^2$; frame rate, 0.33 fps.

**Supplementary Movie 9: Simulation of clustering with particle radius = 13 nm.**
Single units are shown as gray circles; cluster members are shown as colored circles, with identical colors indicating membership of the same cluster. Frames were taken every 5 s over a 750-s simulation. Field of view, 500 × 500 $nm^2$; frame rate, 10 fps.

**Supplementary Movie 10: Simulation of clustering with particle radius = 16 nm.**
Single units are shown as gray circles; cluster members are shown as colored circles, with identical colors indicating membership of the same cluster. Frames were taken every 5 s over a 750-s simulation. Field of view, 500 × 500 $nm^2$; frame rate, 10 fps.

**Supplementary Movie 11: Simulation of clustering with 13 fixed particles (particle radius = 13 nm).**
Red dashed circles highlight fixed particles. Single units are shown as gray circles; cluster members are shown as colored circles, with identical colors indicating membership of the same cluster. Frames were taken every 5 s over a 750-s simulation. Field of view, 500 × 500 $nm^2$; frame rate, 10 fps.

**Supplementary Movie 12: Simulation of clustering with 13 fixed particles (particle radius = 16 nm).**
Red dashed circles highlight fixed particles. Single units are shown as gray circles; cluster members are shown as colored circles, with identical colors indicating membership of the same cluster. Frames were taken every 5 s over a 750-s simulation. Field of view, 500 × 500 $nm^2$; frame rate, 10 fps.
